# More fungi than legs: the first fungal microbiome for a fungus-eating millipede (Colobognatha)

**DOI:** 10.1101/2024.12.23.629787

**Authors:** Angie Macias, Brian Lovett, Michelle Jusino, Lauren Cole, Matt T. Kasson

## Abstract

Fungi are widely consumed across the animal kingdom for nutritional and defensive benefits. Millipedes, among the first air-breathing land animals, were also among the first terrestrial fungivores. As detritus-eating omnivores, most millipedes regularly consume fungi. Millipede diets diverged ∼200 million years ago, when obligate fungivorous millipedes (subterclass Colobognatha) diverged from their detritivorous counterparts. Despite their global distribution and uncommon diet, little is known about the association between Colobognaths and the fungi they consume. In 2019, surveys of fungal communities associated with the Colobognath *Brachycybe lecontii* revealed associations with at least 176 genera of culturable fungi. Given the known biases of culture-based approaches, a more comprehensive survey of *B. lecontii*’s fungal microbiome using amplicon sequencing was undertaken. In this study, we generated amplicon sequence data to look for associations between fungi and *B. lecontii*, and to determine if patterns of fungal diversity are millipede- or habitat-driven. Altogether, the fungal microbiome of *B. lecontii* encompassed 620 fungal genera and 100 orders. Despite much greater observed fungal diversity in the amplicon-based study, sampling was likely not sufficient to capture the full diversity of fungi associated with *B. lecontii*. Taxonomic and functional diversity were significantly influenced by site and colony, indicating that community structure is shaped by geography and habitat. It remains unknown whether these findings are representative of the larger patterns of fungal diversity for the entire millipede subterclass. Nevertheless, the obligate fungivorous lifestyle employed by these long-extant animals may provide important clues regarding fungal diversity and function.

## Introduction

Many animals depend on fungi for food, water, defense, and shelter. Mycophagy (the consumption of fungi), especially facultative mycophagy, is both widespread and common among animals (Claridge and Trappe 2005). Through consumption, animals can indirectly access recalcitrant resources due to the enzymatic & bioaccumulative abilities of fungi (Douglas 2022, Byzov 2006, Birkemoe et al. 2018). Nutritionally, fungi are generally rich in fiber, amino acids, protein, carbohydrates, phosphorus, and potassium, and are a good source of the trace elements manganese, copper, zinc, and especially selenium (Claridge and Trappe 2005, Martin 1979). Due to their high (>80%) water content, fungi are also a valuable resource in arid environments (Claridge and Trappe 2005, Elliott et al. 2022).

Mycophagy has been reported in mammals (Elliott et al. 2022), birds (Elliott et al. 2019b), reptiles (Elliott et al. 2019a), amphibians (Rossa-Feres et al. 2004), fish (Neal et al. 2020), insects (Hanski 1989, Douglas 2022), collembolans (Hammond and Lawrence 1989), mollusks (Silliman and Newell 2003, Ori et al. 2021), crustaceans (Thomas and Thomas 2022), arachnids (Ruess and Lussenhop 2005, Santamaria et al. 2023), nematodes (Ruess and Lussenhop 2005, Zhang et al. 2020), and millipedes (Macias et al. 2019). The most well-known examples of obligate mycophagy include “fungus farming” leafcutter ants (Mueller et al. 2005), ambrosia beetles (Beaver 1989, Hulcr and Stelinski 2017), termites (Mueller et al. 2005), and *Sirex* wood wasps (Coutts and Dolezal 1969). More recently discovered examples of obligate mycophagy among invertebrates include mushroom-harvesting ants (von Beeren et al. 2014), lizard beetles (Toki et al. 2012), marsh periwinkle snails (Silliman and Newell 2003), and Brazilian stingless bees (Menezes et al. 2015). Among vertebrates, obligate mycophagy has been reported from red-backed voles (Maser and Maser 1988, Claridge and Trappe 2005), potoroos (Sinclair et al. 1996, Green et al. 1999, Claridge and Trappe 2005), bettongs (Elliott et al. 2022), northern flying squirrels (Lehmkuhl et al. 2004), and parrots (Elliott et al. 2019b).

In addition to nutritional benefits, some animals use fungi or their secondary metabolites in defensive mutualisms. These partnerships involve a fungal mutualist improving a host’s resistance to outside antagonists, either through improving host vigor and/or immunity, through competitively excluding pathogens and other threats, or through producing secondary metabolites (Flórez et al. 2015, Douglas 2022). Microbial defensive symbioses were recently reviewed by Flórez et al. (2015), who summarized relationships between fungi and ants (Rodrigues et al. 2009, Carriero et al. 2002, Van Bael et al. 2009, Wang et al. 1999, Hervey and Nair 1979), beetles (Davis et al. 2011, Nakashima et al. 1982, Guo et al. 2023), termites (Aanen et al. 2009), fruit flies (Rohlfs and Kürschner 2010), and honeybees (Gilliam et al. 1988). More recent studies have revealed the defensive roles of fungi in several mutualisms, including the anti-nematode activity of *Amylostereum* in wood wasps (Hajek et al. 2019), egg protection by Cordycipitaceae fungi in stinkbugs (Nishino et al. 2024), and the food preserving abilities of *Yarrowia* yeasts in burying beetles (Shukla et al. 2018, Biedermann and Vega 2020). In most cases, the presence, identity, or specific activity of protective secondary metabolites have yet to be formally investigated.

Millipedes have long been known to associate with and consume fungi (Hopkin and Read 1992), yet the identity of the fungi involved and the specific benefits that millipedes receive are largely unexplored. Most extant millipedes are omnivorous detritivores that regularly consume fungi as part of their natural diet. The fungal associations of 16 millipede species from seven families have been investigated but their level of dependence on these fungi is unknown (Table 1, Supplemental Table 1). Obligate mycophagy has been identified in the Colobognatha, a subterclass with ∼300 formally described species across four orders (MilliBase.org). With the exception of our previous culture-based investigations of *Brachycybe lecontii* (Macias et al. 2019), the fungal communities of Colobognaths have never been formally investigated. This subterclass was the most dominant millipede group 100 million years ago according to the fossil record (Wesener and Moritz 2018), but today represents just over 2% of known millipede species. These obligate fungivores diverged from their detritivorous counterparts >200 million years ago, and as such, are among the oldest obligate fungus-feeding animals on the planet. These millipedes lack robust mandibles and associated musculature needed for chewing (Hopkin and Read 1992) and instead have either suctorial mouthparts (Moritz et al. 2022) or mouthparts adapted for a scraping–slurping mode of feeding (Moritz et al. 2021, Wong et al. 2020).

**TABLE 1.**
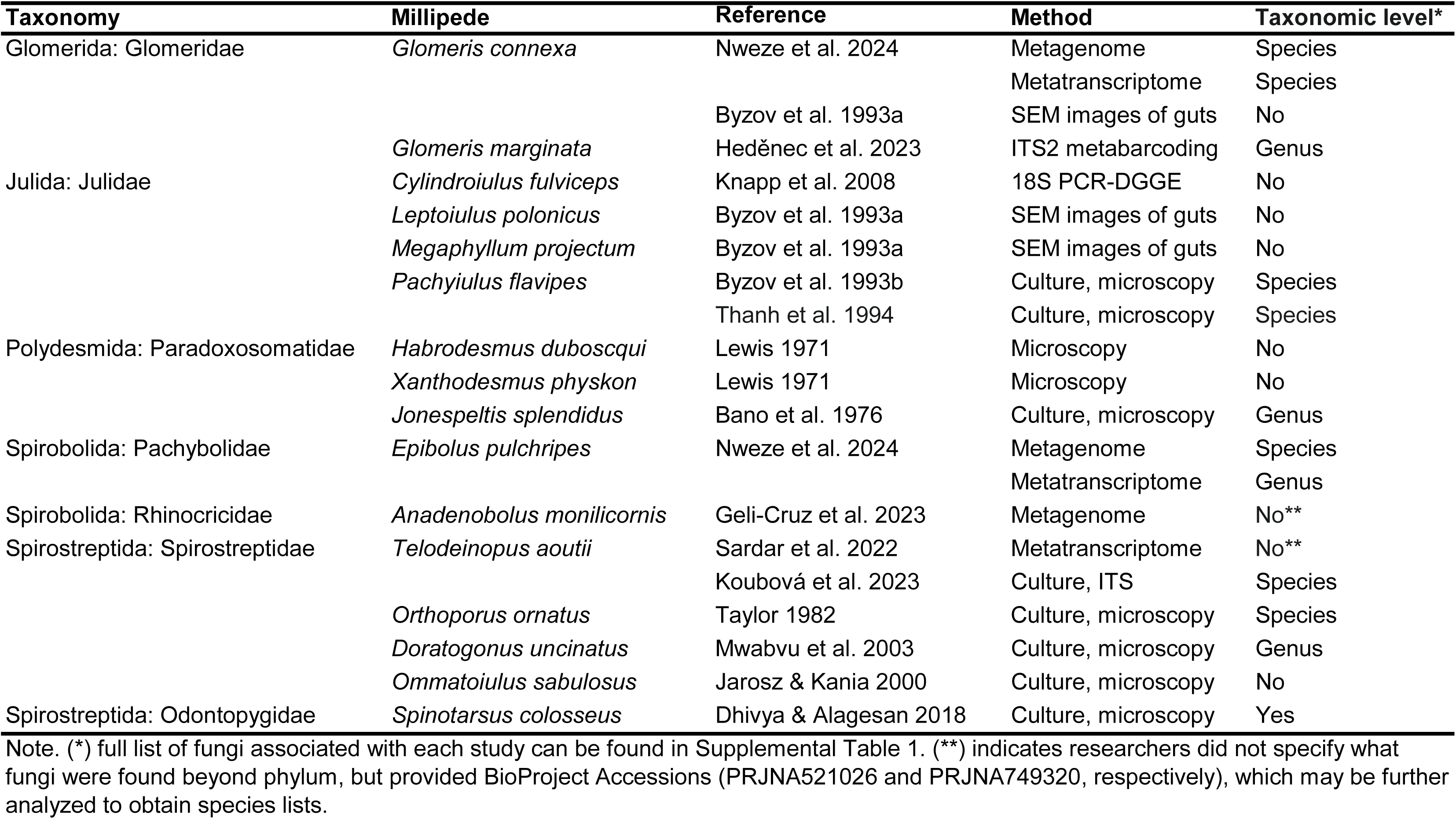
List of all studies that found associations between millipedes and fungi.

Colobognath millipedes are severely understudied relative to other invertebrate arthropods, despite their global distribution and uncommon diet (Read and Enghoff 2009, Shorter et al. 2018). As of November 2024, The International Union for Conservation of Nature’s Red List of Threatened Species has assessed 250 millipedes, which represents just 1.9% of known millipede species (IUCN 2024). Of the seven Colobognath millipedes assessed, 2 are vulnerable, 4 are endangered, and 1 is critically endangered. The U.S. Fish and Wildlife Service, who is responsible for the same classifications in the United States, has not assessed the conservation status of any millipedes, including the 35 Colobognatha millipedes in the U.S. (USFWS ECOS, 2024).

Recent research on Colobognaths has focused on the charismatic genus *Brachycybe*, commonly referred to as feather millipedes. This includes studies on their anatomy (Blanke and Wesener 2014, Moritz et al. 2021), evolution (Brewer et al. 2012), fungus-based diet (Macias et al. 2019), natural history (Wong et al. 2020) and chemical defenses (Jones et al. 2022, Banks et al. 2024). *Brachycybe* includes eight named species and at least 2 undescribed species, with most species in the United States, and others in China, Taiwan, South Korea, and Japan (Brewer et al. 2012). One species, *B. lecontii*, is known to associate with at least 176 genera of culturable fungi in 39 orders and has been observed directly feeding on various fungus-colonized substrates in the field and on pure fungal cultures in the lab (Macias et al. 2019, Wong et al. 2020). Our 2019 culture-based study was the first survey of fungal communities associated with any Colobognath millipede (Macias et al. 2019). Wong et al. (2020) provided additional insight into fungus feeding by *B. lecontii*, highlighting bowl-shaped wet depressions that often appear in fungal tissues underneath feeding millipedes and the anatomic features necessary for such feeding including a labral brush and tarsal combs. *B. lecontii* also possess large cephalic glands that may be involved in saliva production, which may aid in external digestion of fungi (Blanke and Wesener 2014). The lack of solid matter in dissected digestive tracts of *B. lecontii* coupled with clear liquid regurgitation and defecation observed in this species offers strong support for liquid-based fungus-derived diet (Wong et al. 2020).

*Brachycybe lecontii* has four distinct evolutionary lineages with some degree of geographic separation (Brewer et al. 2012), but insufficient sequence data was available to adequately resolve these clades. *Brachycybe* are subsocial, with groups of several to 100+ individuals forming stable persistent stellate aggregations (pinwheels) while feeding on fungi (Wong et al. 2020). Males also protect eggs by grooming them and transporting them away from danger and abstain from feeding until the young hatch (Wong et al. 2020, Murakami 1962, Kudo et al. 2010). This exclusive paternal care, a rare phenomenon among animals, is required for breeding success (Murakami 1962, Kudo et al. 2010).

Another notable aspect of *Brachycybe* biology is the production of defensive alkaloids, including deoxybuzonamine, gosodesmine, hydrogeosodesmine, homo-geosodesmine, and homo-hydrogeosodesmine (Jones et al. 2022, Banks et al. 2024). Buzonamine, a millipede compound related to deoxybuzonamine, was found to repel predatory ants (Wood et al. 2000), but any anti-predatory effects of the compounds found in *Brachycybe* remains unknown, as do any anti-fungal, anti-bacterial, or anti-parasitic effects. However, these millipedes only produce visible droplets of the compound when physically disturbed, supporting a defensive role (Wong et al. 2020, Banks et al. 2024). While it is believed that *B. lecontii* synthesize their defensive compounds *de novo*, pyrrolidine alkaloids similar to those produced by *B. lecontii* are known from several fungi including *Aspergillus* spp. (Xu et al. 2020), *Epichloë* spp. (Pan et al. 2014), and *Humicola* sp. (Gomi et al. 1993).

While the culturable fungal community associated with *B. lecontii* has been described (Macias et al. 2019), the conclusions of this study were limited by culture biases and sample size variability. Diversity metrics were linearly related to sample size, and 40% of all isolates were the sole representatives for their genus, despite sampling over 300 millipedes over several years (Macias et al. 2019). In this current study, we collected millipedes from three sites along the Appalachian Mountains, which are the center of origin for *B. lecontii* (Brewer et al. 2012). We included two of the same sites from the Macias et al. 2019 study, allowing for direct comparison between the two studies. Altogether, we sought to answer four main questions: 1) What is the composition of the fungal microbiome associated with *B. lecontii*, as identified with amplicon sequencing?; 2) Is fungal diversity (compositional, functional, and abundance) driven by millipede sex, colony, or collection site?; 3) How does the fungal microbiome of *B. lecontii* compare to our previous culture-based study and fungal communities of non-Colobognath millipedes?; and 4) Is there any evidence to-date for fungal-millipede mutualisms? Answering these questions will lead to a better understanding of the factors shaping the fungal microbiome of *B. lecontii* and other fungus-feeding millipedes and will allow a deeper understanding of arthropod-fungal interactions.

## Methods

### Collections

Five sites were chosen to create a northeast-southwest transect along the Appalachian Mountains, covering the northeastern portion of *B. lecontii*’s known distribution (Figure 1). Of the five surveyed sites, *B. lecontii* was recovered in three sites: Cove Lake State Park (abbreviated as CO) in Caryville, Tennessee (36.307, -84.227); Bad Branch State Nature Preserve (BA) in Eolia, Kentucky (37.068, -82.772); and Chief Logan State Park (CH) in Logan, West Virginia (37.893, -82.013). *B. lecontii* were never recovered from two of five surveyed sites: Kentucky Ridge State Forest (KR) in Pineville, Kentucky (36.693, -83.797), and Breaks Interstate Park (BI) in Elkhorn City, Kentucky (37.301, -82.311). Site CO is a damp, east-facing slope dominated by beech, maple, and hemlock, and was sampled in a previous culture-based study in 2015 (TN1; Macias et al. 2019). Site BA is a dry valley downstream of a hemlock and rhododendron forest, and is dominated by tulip poplar, white oak, and maple. Site CH is a dry northeast-facing slope dominated by maple, tulip poplar, and black walnut, and was sampled in a previous culture-based study in 2016 (WV5; Macias et al. 2019). Sampling was conducted during August 16-18, 2019.

**Figure 1:**
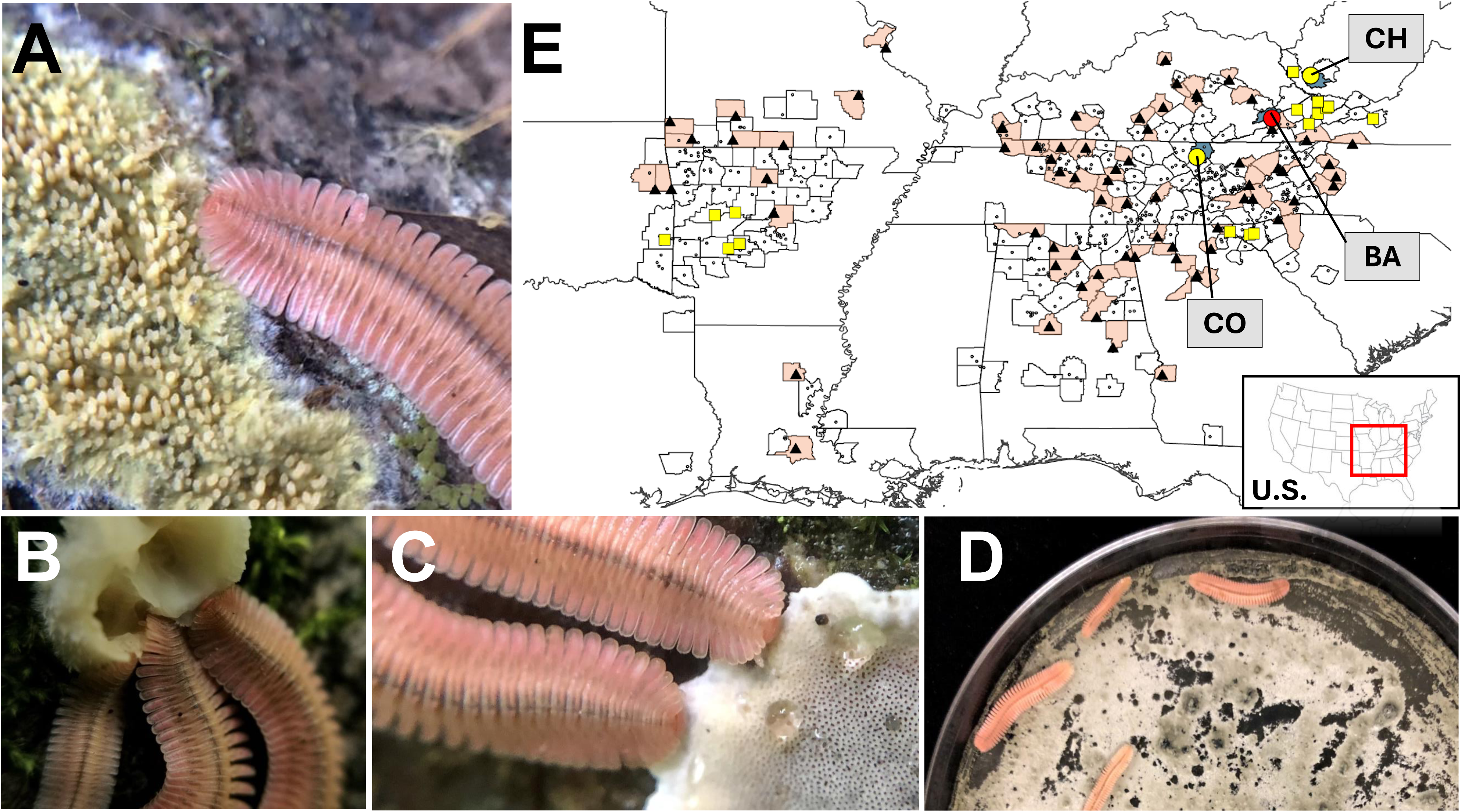
*Brachycybe lecontii* feeding on fungi (A-D) and a range map for the species (E), with observations from the literature, iNaturalist, and BugGuide in black dots, records from Macias et al. 2019 in yellow squares (includes CO and CH in yellow circles), and records from this study in red circles. All drawn counties have *B. lecontii* records, but counties shaded red are new records not yet in the literature. Blue counties were sampled in this study. Inset map shows U.S. range footprint. Photos A-C taken in situ at site CH in 2019 and 2020. Photo D shows field-collected *B. lecontii* feeding on a pure culture of *Oidiodeondron* sp. BC551 from site CO.

Millipedes were located by turning over rotting woody debris in habitats appropriate for *B. lecontii*. Only mature adult millipedes were collected, and where possible, we attempted to have equal numbers of males and females and equal-sized samples from different colonies. Information about individual samples can be found in Supplemental Table 2. Millipedes were sexed using a hand lens (Wong et al. 2020), and juveniles or excess adults were returned to their colony. Logs were returned to their initial position after sampling. Millipedes were collected with flame-sterilized soft forceps and placed into sterile 2 mL screw-top tubes pre-filled with 95% ethanol. These tubes were held on ice for up to 2 days in transit, and stored in a -80 freezer until extraction. The map in Figure 1 was generated using ArcGIS® software by Esri. ArcGIS® and ArcMap™ are the intellectual property of Esri and are used herein under license.

### DNA extraction

Whole millipedes were dried, weighed, and photographed in a sterile hood prior to DNA extraction. A modified DNeasy PowerSoil Pro kit (Qiagen: Germantown, MD, USA) was used for DNA extraction. Vortexing with glass beads was insufficient to break up the large amount of exoskeleton, so millipedes were macerated by hand using a DNA-sterile micropestle in a 1.5 mL DNA-sterile tube, then vortexed according to the protocol. DNA quantity was estimated using a Qubit 3.0 Fluorometer (Thermo Fisher Scientific: Waltham, MA, USA) and a Qubit dsDNA HS Quantification Assay Kit (Thermo Fisher Scientific). Millipedes that yielded too little DNA were omitted from downstream processing.

For controls, DNA extractions of lysis buffer, a single-copy synthetic mock community containing ITS-like sequences (SynMock; Palmer et al. 2018), and a mock biological community (BioMock) were used. The DNA extractions of lysis buffer serve as negative controls, allowing filtering of taxa that likely originate as reagent contamination. SynMock was used to filter sequences to reduce the effects of tag-switching and to parameterize bioinformatic processing (Palmer et al. 2018). A BioMock community was generated to detect biases in amplification or sequencing depth by taxon, and to assist in detecting important missing data in downstream analyses (Palmer et al. 2018). Our BioMock consisted of pooled equilibrated DNA aliquots extracted from pure cultures of the twelve fungi found to be central to the *Brachycybe lecontii* culturable fungal community (Macias et al. 2019), in addition to one undescribed genus from the same study: *Phialophora* (Ascomycota: Chaetothyriales), *Fonsecaea* (Ascomycota: Chaetothyriales), *Ramichloridium* (Ascomycota: Mycosphaerellales), *Cosmospora* (Ascomycota: Hypocreales), *Verticillium* (Ascomycota: Hypocreales), *Trichoderma* (Ascomycota: Hypocreales), *Penicillium* (Ascomycota: Eurotiales), *Phlebia* (Basidiomycota: Polyporales), *Irpex* (Basidiomycota: Polyporales), aff. *Apophysomyces* (Mucoromycota: Mucorales), *Mucor* (Mucoromycota: Mucorales), *Umbelopsis* (Mucoromycota: Umbelopsidales), and *Actinomortierella* (Mucoromycota: Mortierellales). Fungal DNA was extracted and quantified as above.

### Primers, library prep, & sequencing

The universal fungal barcoding locus, the nuclear ribosomal internal transcribed spacer region (ITS), was chosen for this study, as it is most well-represented in sequence databases and offers better taxonomic resolution (Schoch et al. 2012). The full-length ITS locus is too long for the chosen sequencing method, however, so the ITS2 sub-region was used. ITS2 (and full-length ITS) is length- and sequence-variable and can have multiple paralogous or non-orthologous copies throughout the genome (Schoch et al. 2012). The ITS locus also sometimes has mutations in popular priming sites that can impede sequencing (Reynolds et al. 2022). Early diverging fungi in particular, such as the Zoopagomycota, often fail to amplify due to priming site mutations and extended length (Reynolds et al. 2019, 2022). Altogether, these factors can impede sequence amplification and OTU identification, or prevent certain analyses based on read counts or phylogeny.

Dual-indexing library prep, a half-lane of Illumina MiSeq v3 2×300 sequencing, and FastQC analysis were conducted at the University of Minnesota’s Genomics Core Facility (UMGC), using the ITS2 primer pair 5.8SR (TCGATGAAGAACGCAGCG; Hopple and Vilgalys 1999) & ITS4 (TCCTCCGCTTATTGATATGC; White et al. 1990), both modified with Illumina Nextera adapters. UMGC reports that all PCR steps use KAPA HiFi Hot Start Polymerase (Roche Sequencing Solutions: Pleasanton, CA, USA), and use the above primer pair, except for PCR2, which uses a mix of forward and reverse indexing primers. Initial qPCR was conducted using the following conditions: initial denaturation at 95°C for 5 minutes, 35 cycles of denaturation at 98°C for 20 seconds, annealing at 55°C for 15 seconds, and extension at 72°C for 60 seconds, followed by final extension at 72°C for 5 minutes. After qPCR, samples were normalized to 167,000 molecules/μL. PCR1 used the same conditions, except only 25 cycles, and PCR2 as well, except only 10 cycles. Pooled sample was denatured with NaOH, diluted to 8 pM in Illumina’s HT1 buffer (Illumina: San Diego, CA, USA), spiked with 15% PhiX, and denatured at 96°C for 2 minutes immediately before loading. Raw sequencing data is available at NCBI’s Sequence Read Archive (BioProject ###).

### Bioinformatics

Demultiplexed sequencing data was processed with the AMPtk pipeline v. 1.5.2 (Palmer et al. 2018), with additional steps using LULU (Frøslev et al. 2017), BASTA v. 1.4.0 (Kahlke and Ralph 2018), NCBI BLAST+ (Camacho et al. 2009), and R v. 4.0.3 (R Core Team 2020). AMPtk was chosen over other pipelines due to its ability to analyze variable-length amplicons and to filter based on spike-in controls. Up to two nucleotide mismatches were allowed in each primer, a maximum of one mismatch allowed in the barcode, the maximum length was set to 300 bp (longer reads are truncated), an index bleed rate of 1.8% was detected and filtered, and reference-based chimera filtering was used during clustering. A minimum length of 150 bp was used, and the clustering threshold was 97% similarity using the DADA2 algorithm (Callahan et al. 2016). For taxonomy assignment, two major steps were used, 1) AMPtk’s hybrid method using the built-in UNITE database and SINTAX, followed by 2) re-assignment for OTUs not resolved to class or better, using NCBI BLAST+ and BASTA in August 2021. OTUs found in the negative control samples were dropped if 10% or more of all reads for the OTU were from the negative controls. OTU tables were trimmed to remove samples with <5,000 reads, and LULU was used to merge OTUs identified as artifacts without discarding rare but real OTUs. Code is available in Supplemental File 1 & 2.

### Statistical analyses

Once the OTU table was obtained, further analyses were conducted in R, with code available in Supplemental File 2. To estimate if sampling captured a representative view of the true fungal diversity, a taxon accumulation plot (Fisher et al. 1943) was generated using the function specaccum in the package vegan v. 2.5.7 (Oksanen et al. 2020). Shannon diversity index (SDI; Shannon 1948), a metric of alpha diversity, was calculated using the diversity function in vegan. Only millipede sexes and sampling sites could be fully statistically compared, while only select colonies could be compared. Shapiro-Wilk normality test (function shapiro.test in stats 4.0.3; R Core Team 2020; Shapiro & Wilk, 1965) was used to assess normality assumptions, and the Bartlett test of homogeneity of variances (function bartlett.test in stats; Bartlett 1937) or Levene’s test (levene_test in rstatix v. 0.7.0; Kassambara 2023; Levene 1960) were used to test equality of variance assumptions. The functions named below are all in package rstatix. A Welch’s ANOVA (welch_anova_test; Welch 1951) and a Games-Howell posthoc test (games_howell_test; Games & Howell 1976) were used to compare sites, and a standard t-test (t_test) was used to compare sexes. At the colony level, two comparisons were assessed: two large colonies of similar size from the same site (Wilcoxon Test, using wilcox_test; Wilcoxon 1945); and all colonies with at least four individuals sampled (ANOVA, using anova_test; Fisher 1918). To assess if millipede dry weight, millipede sex, and SDI were correlated, Pearson’s product moment correlation coefficient (Pearson 1895) was calculated using the function cor.test in R package stats.

For beta diversity, dissimilarity matrices were generated with the function vegdist in vegan, using Bray-Curtis dissimilarity (abundance-based), and with the function betadiver using βsim dissimilarity (presence / absence-based). To compare the effects of collection site, millipede sex, and millipede colony on fungal community composition, permutational multivariate analysis of variance (PERMANOVA; function adonis2 in vegan; Anderson 2008) was conducted, using pairwise comparisons when appropriate (function pairwise.adonis in package pairwiseAdonis v. 0.4; Arbizu 2017). Effect size (⍵^2^, R^2^) is given for significant comparisons, and strength assessment benchmarks from the R package effectsize v. 0.8.9 (Shachar et al. 2020) were used for interpretation.

The Bray-Curtis & βsim dissimilarity matrices were used to generate Principal Coordinate Analysis plots (PCoA; function cmdscale in stats; Gower 1966), and Nonmetric Multidimensional Scaling (NMDS; function metaMDS in vegan; Shepard 1962a, 1962b) plots using ggplot2 v. 3.4.4 (Wickham 2016). A stress plot was also generated for the NMDS using the function stressplot in vegan. NMDS analyses were validated using the permutation-based method described in Dexter et al. 2018. The PCoA axes reflected very little of the variation in the dataset, therefore only the NMDS will be referred to in the body of this text. The PCoAs are available in Supplemental Figures 1 and 2.

### Culture-based dataset re-analysis

To compare results from Macias et al. 2019 to this study, a re-analysis of the original dataset was required. The original Sanger sequences were re-cleaned and re-trimmed by examining the original chromatograms by eye and correcting mis-called or ambiguous bases. Any sample missing important metadata (such as the millipede from which it was collected) was dropped, leaving 826 sequences. Since these sequences resulted from isolations of single fungal strains, clustering was not used. Instead, these cleaned Sanger sequences were fed directly into the AMPtk Taxonomy command, to ensure that the culture-free and culture-based datasets would be comparable. BLAST+ and BASTA were also used as described above, except searches were conducted in April 2022. Assigned taxonomy and metadata are available in Supplemental Table 3, and code is available in Supplemental File 1.

### Functional analysis

To assess the functional diversity of communities recovered from the culture-based dataset and this study, the functional trait databases FunGuild (Nguyen et al. 2016) and FungalTraits (Pӧlme et al. 2021) were queried. For simplicity, only results from FunGuild will be referred to in the body of this text, but the FungalTraits results are available in Supplemental Tables 4 and 5. Some FunGuild guilds were merged to simplify analysis (Supplemental File 2).

To compare the functional diversity of fungal communities across sites, colonies, and sexes in this study, PERMANOVA (adonis2) and pairwise PERMANOVA (pairwise.adonis) tests were conducted on Bray-Curtis distance matrices using only OTUs that were successfully assigned to a guild using FunGuild. An NMDS (metaMDS) plot and companion stress plot (stressplot) were also generated to visualize differences in guild across groups. Code is available in Supplemental File 2.

## Results

### Occurrence, collections, & DNA extraction

As a result of compiling occurrence records for Figure 1, we found 73 unpublished new county records for *B. lecontii* in 11 of 15 states where it is found, a 30% increase in the known range for the species. The great majority of these records came from the community science website iNaturalist. This data search did not examine museum specimens for which there is no associated publication, or museum records for which no data is available online.

Sixty-eight millipedes were collected for the culture-free study, and 55 remained in the final dataset after quality control steps (Supplemental Table 2). At site CO, 20 millipedes were collected from 5 colonies, all found on downed maple or tulip-poplar logs / branches with visible fungus or lichen. At site BA, 20 millipedes were collected from 2 colonies, all found on downed birch or tulip-poplar logs / branches with little visible fungus. At site CH, 28 millipedes were collected from 8 colonies found on downed maple, beech, tree-of-heaven, or tulip-poplar logs / branches with or without visible fungus. Across all sites, millipedes weighed 23 mg (dry) on average, or 27 mg for females and 18 mg for males. Millipede dry weight was not correlated with DNA yield (p = 0.39): males yielded a mean of 10.0 ng/uL DNA (median 3.81 ng/uL), and females 10.9 ng/uL (median 10.1 ng/uL).

### Sequencing

The culture free dataset included 60 millipedes, 3 negative controls, SynMock, and 2 replicate Biomocks, for a total of 66 samples. Illumina sequencing resulted in 7.4M reads and 6,058 OTUs, which was reduced to 5.9M reads and 2,982 OTUs following quality control measures (Supplemental Table 6). 4.99% of reads were dropped due to primer incompatibility, and 0.04% dropped due to short length. Forward and reverse reads were successfully paired, and read counts were not normalized across samples. The mean number of reads per sample after quality control steps was 98K (5.8K – 161K). A taxon accumulation curve was generated (Supplemental Figure 3). Following taxonomy assignment in AMPtk, 79 (1.3%) OTUs were unassigned, 370 (12.4%) to only kingdom Fungi, and 318 (10.7%) to only phylum. Follow-up analysis of these OTUs with BLAST and BASTA successfully assigned class-or-better taxonomy to 66 (8.7%) of these OTUs.

Of all OTUs generated, 2,863 (96%) were fungal: other kingdoms detected were Viridiplantae (68 OTUs), Alveolata (5), and Metazoa (4). The majority of fungal OTUs came from phylum Ascomycota (1,874, 65.5%), followed by Basidiomycota (611, 21.3%). The proportions of all detected phyla are given in Table 2. Dominant classes in the Ascomycota were Dothideomycetes (1.3M reads, 23.6%) and Sordariomycetes (1.2M reads, 22%), while dominant Basidiomycota classes were Agaricomycetes (0.66M reads, 11.8%) and Tremellomycetes (0.31M reads, 5.5%) (Figure 2). Across the entire study, the most common orders were Hypocreales (Ascomycota: Sordariomycetes; 0.8M reads, 14.2%), Helotiales (Ascomycota: Leotiomycetes; 0.76M reads, 13.5%), and Capnodiales (Ascomycota: Dothideomycetes; 0.74M reads, 13.1%). Each millipede yielded, on average, 35 fungal orders. In total, 9 phyla, 32 classes, 100 orders, 241 families, 620 genera, and 697 species were identified in this study.

**TABLE 2.** Summary of the proportion of OTUs or reads (from amplicon-based study) or of isolates (culture-based study) from each fungal phylum.

| **Phylum** | **OTUs** |  | **Reads** |  | **Isolates** |  |
| --- | --- | --- | --- | --- | --- | --- |
|  | # | % | # | % | # | % |
| Ascomycota | 1874 | 65.46% | 4269993 | 76.10% | 612 | 74.09% |
| Basidiomycota | 611 | 21.34% | 1015618 | 18.10% | 95 | 11.50% |
| Mucoromycota | 6 | 0.21% | 34515 | 0.62% | 68 | 8.23% |
| Mortierellomycota | 5 | 0.17% | 10179 | 0.18% | 33 | 4.00% |
| Basidiobolomycota | 2 | 0.07% | 223 | <0.01% | 1 | 0.12% |
| Zoopagomycota | 2 | 0.07% | 124 | <0.01% | - | - |
| Chytridiomycota | 2 | 0.07% | 28 | <0.01% | - | - |
| Glomeromycota | 1 | 0.03% | 95 | <0.01% | - | - |
| Rozellomycota | 1 | 0.03% | 69 | <0.01% | - | - |
| Unclassified fungi | 359 | 12.54% | 280443 | 5.00% | 17 | 2.06% |

**Figure 2:**
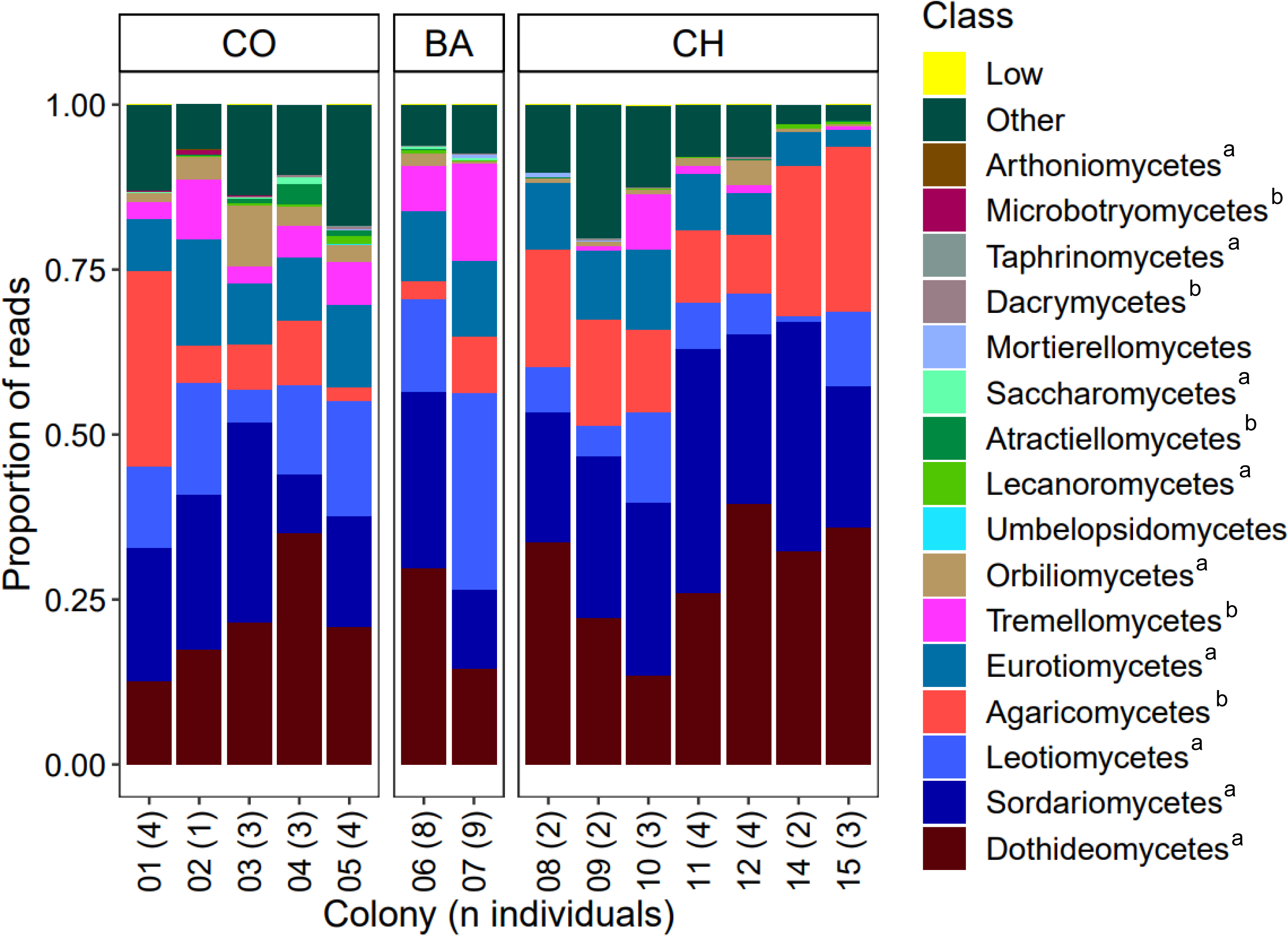
A 100% stacked bar showing the relative proportions of the most common fungal classes in each colony, grouped by site. “Other” = reads identified as “incertae sedis” at the class level or those with no hits. “Low” = 16 classes with fewer than 1,000 reads. Superscripts indicate major fungal phyla: a = Ascomycota, b = Basidiomycota.

There were 6 OTUs found in nearly all (>85%) millipedes: OTU1 *Cladosporium* sp. (absent in 3 millipedes), OTU29 *Scolecobasidium* [*Ochroconis*] *cordanae* (4), OTU2 *Oidiodendron* sp. (5), OTU100 *Helminthosporium asterinum* (6), OTU31 Unknown Mycosphaerellaceae sp. (8), and OTU45 *Cladophialophora* sp. (8). Of these 6 OTUs, 3 have high read counts as well: OTU1 *Cladosporium* sp. (8.5% of all reads), OTU2 *Oidiodendron* sp. (7.1%), and OTU45 *Cladophialophora* sp. (1.7%).

### Culture-based study re-analysis

In the re-analyzed culture-based dataset from Macias et al. 2019, the great majority of isolates came from phylum Ascomycota (612, 74.1%), followed by Basidiomycota (95, 11.5%). The proportions for all phyla are given in Table 2. Dominant classes in the Ascomycota included Sordariomycetes (326 isolates) and Eurotiomycetes (111), while dominant Basidiomycota classes were Agaricomycetes (72) and Tremellomycetes (9) (Supplemental Table 3). Across the entire study, dominant orders were Hypocreales (Ascomycota: Sordariomycetes; 163), Chaetothyriales (Ascomycota: Eurotiomycetes; 72), and Polyporales (Basidiomycota: Agaricomycetes; 57). A total of 93 fungal genera were shared across the culture-free and culture-based datasets. These shared genera included *Cladosporium*, *Oidiodendron* and *Cladophialophora* among others, all of which matched OTUs found in >85% millipedes also with high read counts.

### α and β diversity

Alpha diversity was assessed at the site, colony, sex, and sample level (Table 3) using Shannon’s diversity index (SDI). Site CO had the greatest diversity (5.0), and males had slightly more than females (5.13 and 5.08, respectively). Colonies ranged from 3.7 – 4.7 (mean 4.0), and individuals from 3.0 – 4.5 (mean 3.6). Sites were significantly different (p = 0.04, ⍵^2^= 0.15), specifically CH vs. CO (p = 0.04). Sexes were not significantly different (p = 0.77). Two large colonies from the same site were not significantly different (p = 0.67), and colonies from all sites with four millipedes sampled were also not significantly different (p = 0.60).

**TABLE 3.** Summary of alpha diversity statisitics based on Shannon’s Diversity Index (SDI) for sites (BA, CO, CH), colonies (1-15), individuals (1-32), and sex.

| Site | SDI | Colony | SDI | Sample | SDI |
| --- | --- | --- | --- | --- | --- |
| CO | 4.991 | 1 | 4.344 | 19CO01F | 3.31 |
|  |  |  |  | 19CO02F | 4.063 |
|  |  |  |  | 19CO03F | 2.974 |
|  |  |  |  | 19CO04F | 3.486 |
|  |  | 2 | NA | 19CO06F | 3.609 |
|  |  | 3 | 4.728 | 19CO09M | 4.394 |
|  |  |  |  | 19CO11M | 4.153 |
|  |  |  |  | 19CO12F | 4.539 |
|  |  | 4 | 4.189 | 19CO13F | 4.028 |
|  |  |  |  | 19CO15M | 3.309 |
|  |  |  |  | 19CO16F | 3.978 |
|  |  | 5 | 4.379 | 19CO17F | 3.966 |
|  |  |  |  | 19CO18F | 3.902 |
|  |  |  |  | 19CO19F | 3.841 |
|  |  |  |  | 19CO20M | 4.259 |
| BA | 4.323 | 6 | 4.02 | 19BA01F | 3.2 |
|  |  |  |  | 19BA02M | 3.683 |
|  |  |  |  | 19BA04M | 3.439 |
|  |  |  |  | 19BA05M | 4.321 |
|  |  |  |  | 19BA07F | 3.49 |
|  |  |  |  | 19BA08F | 3.16 |
|  |  |  |  | 19BA09M | 3.211 |
|  |  |  |  | 19BA10F | 3.492 |
|  |  | 7 | 4.044 | 19BA11F | 3.933 |
|  |  |  |  | 19BA12M | 3.207 |
|  |  |  |  | 19BA13M | 3.242 |
|  |  |  |  | 19BA15M | 3.228 |
|  |  |  |  | 19BA16F | 3.621 |
|  |  |  |  | 19BA17F | 3.253 |
|  |  |  |  | 19BA18F | 3.275 |
|  |  |  |  | 19BA19F | 3.907 |
|  |  |  |  | 19BA20F | 4.088 |
| CH | 4.834 | 8 | 3.95 | 19CH01F | 3.623 |
|  |  |  |  | 19CH02M | 3.757 |
|  |  | 9 | 3.715 | 19CH03F | 3.543 |
|  |  |  |  | 19CH04M | 3.325 |
|  |  | 10 | 3.818 | 19CH05M | 3.668 |
|  |  |  |  | 19CH06F | 3.239 |
|  |  |  |  | 19CH09M | 3.36 |
|  |  | 11 | 3.975 | 19CH10M | 3.757 |
|  |  |  |  | 19CH11M | 3.631 |
|  |  |  |  | 19CH13F | 3.602 |
|  |  |  |  | 19CH14F | 3.416 |
|  |  | 12 | 4.187 | 19CH18F | 3.603 |
|  |  |  |  | 19CH30F | 3.869 |
|  |  |  |  | 19CH31F | 3.685 |
|  |  |  |  | 19CH32F | 3.836 |
|  |  | 14 | 3.667 | 19CH23F | 3.421 |
|  |  |  |  | 19CH24M | 3.404 |
|  |  | 15 | 3.671 | 19CH25F | 3.314 |
|  |  |  |  | 19CH27M | 3.255 |
|  |  |  |  | 19CH28M | 3.446 |

Note. (\*) sexes were not significantly different using SDI (Males = 5.128, Females = 5.089), but are reported here. Values represent male vs. female millipede comparisons across the three sites.

Bray-Curtis-based beta diversity is summarized in Supplemental Table 7 and visualized in a NMDS plot in Figure 3, with a companion Shepard stress plot in Supplemental Figure 4. βsim-based beta diversity is summarized in Supplemental Table 8 and visualized in a NMDS plot in Supplemental Figure 5, with a companion Shepard stress plot in Supplemental Figure 6. All sites were found to be significantly different (Bray: overall p < 0.001 and R^2^ = 0.19, pairwise comparisons all p < 0.003. βsim: identical except R^2^ = 0.24), with the greatest distance between sites BA and CH (Bray: 0.696, βsim: 0.572). Sexes were not significantly different (Bray: p = 0.668. βsim: p = 0.915). Two large colonies from the same site were significantly different (Bray: p < 0.001, R^2^ = 0.35. βsim: p < 0.001, R^2^ = 0.317), and colonies from all sites with four millipedes sampled were also significantly different (Bray: p < 0.001, R^2^ = 0.23. βsim: p < 0.001, R^2^ = 0.13).

**Figure 3:**
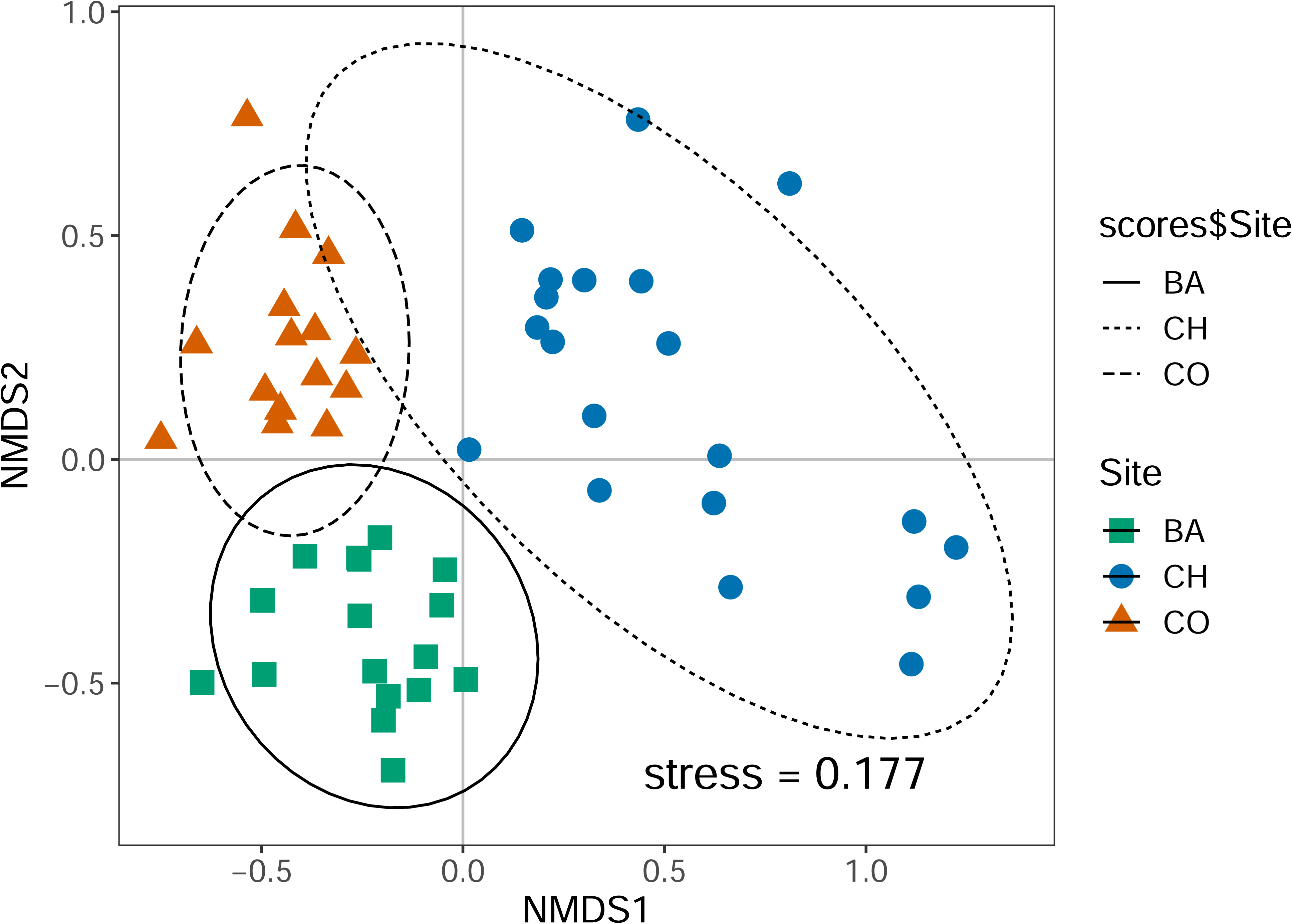
Nonmetric multidimensional scaling (NMDS) ordination plot showing Bray-Curtis dissimilarity scores for all fungi recovered in the culture-free dataset. Point shape & color indicates site: green square = BA, blue circle = CH, orange triangle = CO. Ellipses indicate 95% confidence intervals based on standard error of site centroids. Ellipse line type also indicates site: solid = BA, dotted = CH, dashed = CO. Companion Shepard plot in Supp. Fig. 4.

### Functional analysis

In the culture-based study, FunGuild successfully assigned 98.2% (811) isolates to at least one guild (Supplemental Table 9), while in the culture-free study, only 35.2% (1,009) of OTUs and 45.6% (2.7M) of reads were successfully assigned a guild (Supplemental Table 10). For the culture-based study, 29 guilds were recovered and grouped into ten larger guilds: plant symbiont, plant pathogen, plant parasite, plant saprotroph, epiphyte, saprotroph, animal associate, fungal parasite, lichen parasite, and lichenized. For the culture-free study, 22 guilds were recovered and grouped into the same ten larger guilds as in the culture-based study.

Original group assignments are available in Supplemental Tables 9 and 10, and re-assignments in Supplemental File 2. The functional diversity of fungal communities was compared across sites, colonies, and sexes, and in some cases, across the culture-based and culture-free studies. Differences in the percent of isolates or OTUs assigned to different functional groups in the two studies are shown in Figure 4.

**Figure 4:**
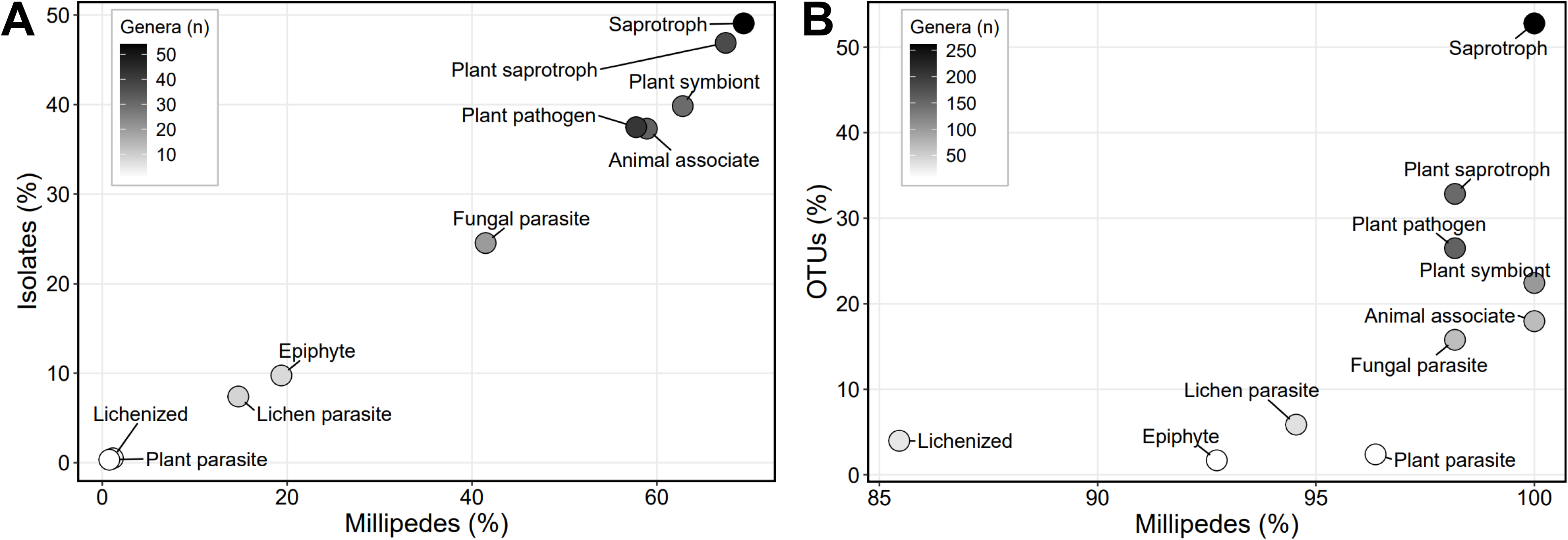
Scatterplots for the culture-based (A) and culture-free (B) studies showing the relative abundance of each guild in terms of the percent of millipedes with 1+ OTUs (A) or isolates (B) in that guild (X-axes) and the percent of OTUs (A) or isolates (B) belonging to that guild (y-axes). Only sequences assigned to a guild are shown. Dot shade represents the number of fungal genera found that fall into the given guild.

In the culture-free dataset, an overall significant difference was found for sites (p = 0.01, R^2^ = 0.1), but pairwise comparisons indicated only BA and CH were significantly different (p = 0.01). Site-level comparisons for culture-based vs. culture-free studies are also summarized in Table 4 and Figure 5 (Shepard plot in Supplemental Figure 7). Millipede sex did not have a significant effect on the functional diversity of fungal communities (p = 0.39). Two large colonies from the same site were significantly different (p < 0.01, R^2^ = 0.26), but colonies from all sites with four millipedes sampled were not significantly different (p = 0.14).

**Figure 5:**
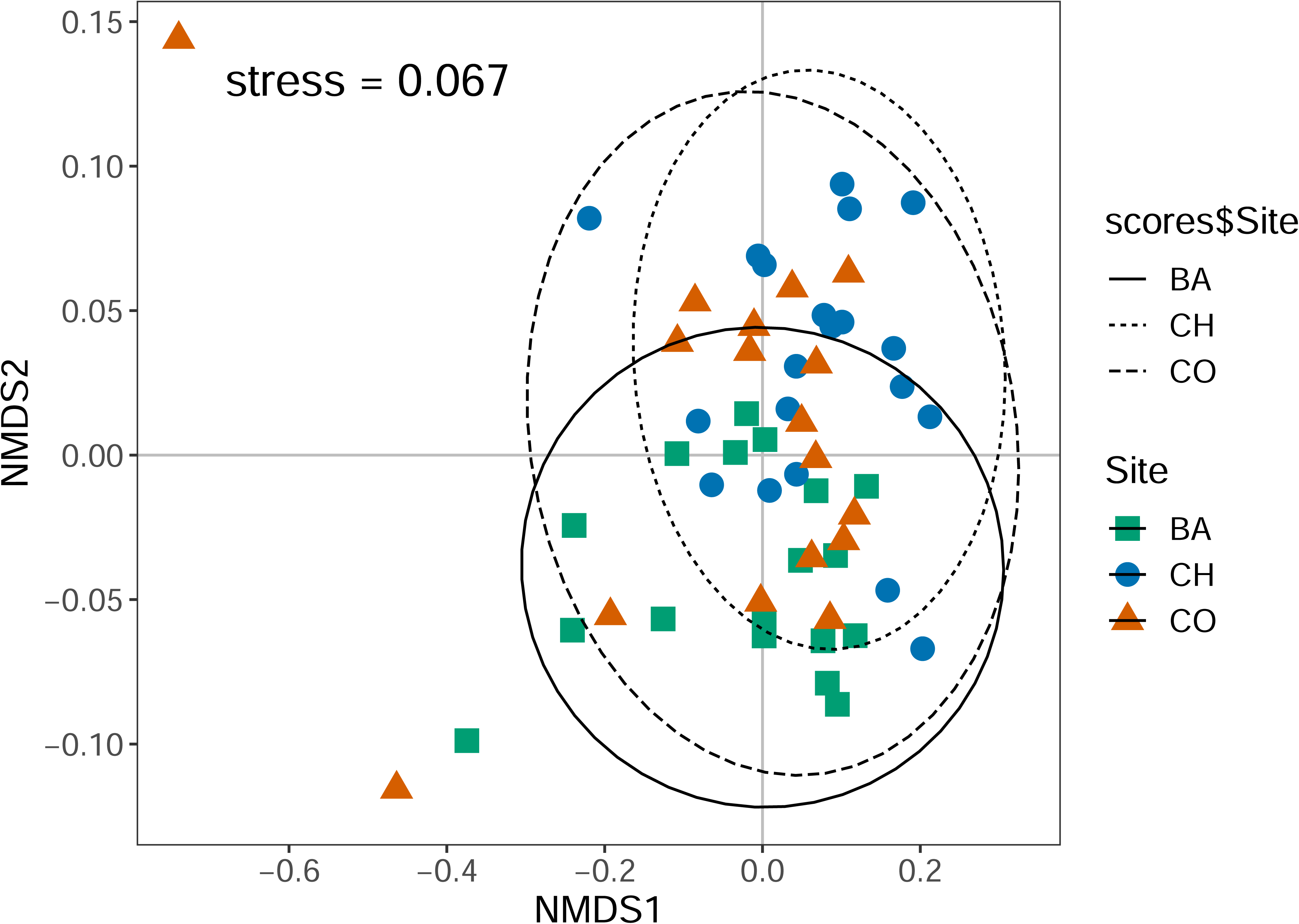
Nonmetric multidimensional scaling (NMDS) ordination plot showing Bray-Curtis dissimilarity scores for all fungi recovered in the culture-free dataset. Reads are grouped by guild. Point shape & color indicates site: green square = BA, blue circle = CH, orange triangle = CO. Ellipses indicate 95% confidence intervals based on standard error of site centroids. Ellipse line type also indicates site: solid = BA, dotted = CH, dashed = CO. Shepard plot for this NMDS is in Supp. Fig. 7.

**TABLE 4.** Summary table of the percent of reads or isolates belonging to each site, by study.

| Guild | Cove Lake (CO) |  | Bad Branch (BA) |  | Chief Logan (CH) |  | Overall |  |
| --- | --- | --- | --- | --- | --- | --- | --- | --- |
|  | isolates (%) | reads (%) | isolates (%) | reads (%) | isolates (%) | reads (%) | isolates (%)* | reads (%) |
| Animal associate | 22.5 | 19.3 | --- | 23.9 | 32.2 | 20.3 | 27.4 | 21.1 |
| Epiphyte | 14.7 | 7.6 | --- | 15.7 | 4.4 | 3.3 | 7.2 | 8.3 |
| Fungal parasite | 24.5 | 16.0 | --- | 22.8 | 22.2 | 17.8 | 18.0 | 18.8 |
| Lichen parasite | 1.0 | 5.3 | --- | 2.7 | 7.8 | 7.7 | 5.4 | 5.5 |
| Lichenized | 0.0 | 0.4 | --- | 0.3 | 0.0 | 0.4 | 0.4 | 0.4 |
| Plant parasite | 2.0 | 1.1 | --- | 2.8 | 0.0 | 1.3 | 0.2 | 1.7 |
| Plant pathogen | 28.4 | 21.5 | --- | 18.6 | 34.4 | 44.1 | 27.5 | 30.0 |
| Plant saprotroph | 43.1 | 31.2 | --- | 22.8 | 46.7 | 41.3 | 34.4 | 32.8 |
| Plant symbiont | 38.2 | 36.7 | --- | 33.6 | 34.4 | 35.3 | 29.2 | 35.2 |
| Saprotroph | 33.3 | 62.1 | --- | 69.3 | 35.6 | 66.6 | 36.0 | 66.1 |

Note. A single OTU or isolate can belong to multiple guilds, so columns do not sum to 100%. Site “Bad Branch” was not sampled for the culture-based study. (\*) indicates multiple culture-based collection sites not listed here were included in the overall percentage.

## Discussion

Who is in *B. lecontii*’s fungal community? In this study, we recovered 620 genera of fungi from 9 phyla and 100 orders in association with 55 mycophagous *B. lecontii* millipedes. The re-analysis of our 2019 culture-based study found 148 genera of fungi from 5 phyla and 32 orders in association with ∼300 millipedes. Despite the smaller sampling effort, the observed fungal diversity was far greater in the culture-free study. Yet, even this diversity is an underestimate: the taxon accumulation curve indicated that sampling was likely not sufficient to capture the full diversity of fungi present in this millipede species (Supplemental Figure 3). Nevertheless, this study, much like the pioneering work on fungal communities in red-cockaded woodpecker excavations, has helped uncover tremendous hidden fungal biodiversity in a system that is difficult to characterize without DNA-based tools (Jusino et al. 2015). While this study was not designed to detect new species of fungi, the 2019 study found evidence for several, including at least one putative new genus (Macias et al. 2019). Altogether, these numbers demonstrate that we are only beginning to understand the scope of the relationships between *B. lecontii* and fungi.

Interestingly, Asomycota were recovered in nearly identical proportions from both studies, despite finding over three times as many genera in the culture-free study (Table 2). Basidiomycota were also found in somewhat similar proportions with higher recovery for the culture-free study, while Mucoromycota and Mortierellomycota were more abundant (and likely over-represented) in the culture-based study (Table 2). These discrepancies are not unexpected, as fast-growing fungi are known to be over-represented in studies involving cultures, and DNA-based studies that use PCR are known to differentially amplify certain groups (Reynolds et al. 2022). The BioMock controls used in this study provide direct evidence of this bias in DNA-based studies: differential amplification indicates that Mortierellomycota and Mucoromycota trend towards under-representation, while Ascomycota and Basidiomycota trend towards over-representation (data not shown).

In the functional diversity assessment, we were unable to assign a guild to a majority of OTUs or reads. Despite the missing data, the relative percentages of isolates or OTUs that belong to certain guilds are remarkably similar, given the very different sampling methods, though a few differences are prominent (Table 4). At the two sites (CO and CH) that were sampled in both studies, animal associates and fungal parasites were more common in the culture-based work, while saprotrophs showed the opposite pattern. These differences may indicate that culturing conditions favor fungal parasite over-representation, to the exclusion of their parasitized counterparts. They may also point to culture-free methods’ ability to capture specialist saprotrophs that do not tolerate standard growth media. A third possible explanation is that a component of the defensive secretions of *B. lecontii*, such as deoxybuzonamine, may be anti-microbial and inhibitory towards certain fungi. Though the antifungal properties of deoxybuzonamine remain untested, similar pyrrolidine alkaloids from *Aspergillus sclerotiicarbonarius* and *A. fumigatus* showed moderate to potent antifungal activities (Petersen et al. 2015, Xu et al. 2020).

Analysis of functional diversity was also impacted by culture biases. Nearly all (98%) isolates from the culture-based study were successfully assigned a guild, while only 35% of OTUs or 46% of reads could be assigned to a guild. Databases like FunGuild or FungalTraits are inherently biased towards organisms that can survive in laboratory conditions, since ecological functions are often validated in controlled lab experiments or in the field. These same biases impact taxonomic databases like UNITE and NCBI Nucleotide, for similar reasons (Lücking et al. 2020). These biases and missing data can conceal important information from a researcher: for example, in this study OTU279 is identified as *Mortierella* sp., but is actually a 100% match to a new undescribed species in *Actinomortierella* (Macias et al. 2019, Vandepol et al. 2020). This undescribed species was identified in a network analysis as one of the 12 core community members for *B. lecontii* (Macias et al. 2019). OTU279 was neither common nor abundant in this study, but was almost certainly under-sequenced due to a primer mismatch and amplicon length (data not shown), a known phenomenon in early-diverging fungi (Reynolds et al. 2022). Regardless of the actual sequencing results, it is important for researchers to place new results in the context of their study topic and search for such gaps.

Additionally, the identity assigned to an individual OTU, such as OTU279, may seem unimportant if one assumes databases are representative of the diversity of fungi. However, 83% of fully-named species are missing from even the broadest sequence database (Schoch et al. 2020), and it is estimated that 92-97% of extant fungal species await naming at all (Hawksworth and Lücking 2017). These known-unknowns and unknown-unknowns could represent a large proportion of reads from a sequencing project, could represent high taxonomic levels, and could have significant ecological roles that are unaccounted for in functional analyses.

### What factors shape patterns of fungal diversity?

Macias et al. (2019) compared the fungal communities associated with *B. lecontii* from different millipede clades, millipede sexes, and ecoregions (as a proxy for site). That study found weak but significant effects for millipede clade and ecoregion, and none for sex. The current study compared the fungal communities across sites, colonies, and millipede sexes. We found a weak to moderate relationship between site and fungal community with every metric tested (alpha, beta, and functional diversity). Even stronger effects were identified for colony, especially when comparing two larger colonies from the same site, in terms of beta diversity and functional diversity. Millipede sex was never a significant factor. Given the limited dispersal ability of these animals (Wong et al. 2020), it is not unexpected to find that spatial factors are important to shaping the fungal community. While millipede sex was not significant, future studies should examine other possible millipede effects, such as age or genetic clade (Brewer et al. 2012).

### How do *B. lecontii*’s fungal associates compare to other millipedes?

The study of fungal associations with millipedes is relatively novel and has exclusively focused on non-colobognath millipedes. Previously published studies use small numbers of animals, often after they were used in an unrelated experiment, except for a few recent -omics-based studies of gut associates. Several studies have used culturing and microscopy or ITS barcoding to identify fungi associated with millipedes (Table 1). Collectively, these studies identified 60 genera, mostly ascomycetes, in association with 6 millipede species from 3 orders (Supplemental Table 1).

Three recent -omics studies assessed the fungi from four millipede species. In the first study, the gut of *Anadenobolus monilicornis* was sampled in Puerto Rico for a metagenomic study, which recovered a very small number of ascomycetes (Table 1; Geli-Cruz et al. 2023). Another study used metatranscriptomics to examine fungi in association with the gut of *Telodeinopus aoutii* from a pet shop in the Czech Republic (Sardar et al. 2022). Overall, only 1.1% of all transcripts were fungal, and of these most were Ascomycota and Mucoromycota, with some Basidiomycota, Chytridiomycota, and Zoopagomycota. As in the previous study, fungi were a small component of the millipede holobiont, and are only discussed at the phylum level. The final study used both metagenomics and metatranscriptomics to identify fungi associated with wild-caught *Glomeris connexa* from the Czech Republic, and a lab colony of *Epibolus pulchripes* from an unknown source (Table 1; Nweze et al. 2024). In each case, Ascomycota represented the great majority of reads, though metatranscriptomics also recovered Blastocladiomycota, Basidiomycota, Mucoromycota, and Chytridiomycota for *G. connexa* and Basidiomycota, Mucoromycota, and Chytridiomycota for *E. pulchripes*.

Only a single study has used metabarcoding to identify the fungi in association with a millipede, in this case the dissected gut of 108 wild-caught *Glomeris marginata* from Denmark (Table 1; Heděnec et al. 2023). At the phylum level, Ascomycota was heavily dominant, followed by Mucoromycota. In total, just 13 genera of fungi were recovered, but most sequences were *Talaromyces* (Ascomycota: Eurotiales), *Trichoderma* (Ascomycota: Hypocreales), or *Rhizopus* (Mucoromycota: Mucorales).

Similarly to all these studies, we also found Ascomycota to be the dominant fungal phylum (65.5% of all OTUs) from *B. lecontii*, though a significant proportion of OTUs were Basidiomycota (21.3%). The great majority of fungal genera found in association with any millipede are unique to that specific study and millipede, or are shared between this study and Macias et al. 2019, the only case where a single species was examined in detail twice. Altogether, the limited information from previous studies suggests that non-Colobognath millipedes do not associate with many fungi, nor do they overlap much in terms of fungal diversity.

Regardless, all of the millipedes discussed above are detritivores with much larger body mass compared to *B. lecontii* and are from the subterclass Eugnatha, sister to the Colobognatha. Moreover, they live in locations very distant from the eastern US, and some studies included animals from pet shops or other artificial environments. Last, our study used whole millipedes, not dissected guts, and as a result includes any fungi adhered to the surface of the exoskeleton or present in other parts of the body. As such, making comparisons across studies is impossible, outside of simple lists of names (Supplemental Table 1).

Most of the previous studies discussed above were searching for gut mutualistic microbes. For *Brachycybe* and other colobognaths, it is unclear if their guts are capable of harboring obligate gut mutualists, despite the tremendous fungal diversity that comprise their microbiome. As of 2015, the anatomy of the guts of 7 Eugnatha millipede orders have been examined, and were found to have fairly standard detritivorous arthropod gut morphology with a three-part alimentary tract, with some areas lined with a membrane and others with cuticle (Fontanetti et al. 2015). However, the findings of Moritz et al. (2021, 2022) show that at least the front of the gut (the head and mouth) of several Colobognath species are highly adapted to a fluid diet, and very different from Eugnaths. Future studies of *B. lecontii* or other Colobognath millipedes should obtain baseline gut morphology data for the subterclass, and should attempt to extract these guts for sequencing and imaging to search for resident gut microbes.

### Is there any evidence for fungus-millipede mutualisms?

Based on results from this study and Macias et al. (2019), *Brachycybe lecontii* are likely generalist fungal feeders, with different diets depending on where they live. Our finding that fungal communities differed across sites does not support the idea that *B. lecontii* has a consortium of conserved mutualistic fungi, but the possibility of specific mutualists still remains. However, other studies of core microbiomes have identified several alternative hypotheses that should be considered (Neu et al. 2021). First, microbial community structure may only be discernible in specific millipede sub-populations, or higher fungal taxonomic levels. Second, sampling may be insufficient to capture the microbial community, and thus, identify the core microbiome. Our samples came from a limited portion of *B. lecontii*’s range and captured only a single time point, meaning that the geographic and temporal range studied was limited and may not be representative.

Three OTUs were exceedingly common in most millipedes: *Cladosporium* (OTU1), *Oidiodendron* (OTU2), and *Cladophialophora* (OTU45). The 2019 study also found several isolates with >99.5% sequence similarity to these specific OTUs. Though these genera were neither common nor abundant in the 2019 study, they were recovered from the same geographic area as the current study. The prevalence and abundance of these fungi in this culture-free study suggest that these three taxa may be significant, but they may be rarer in the larger range of *B. lecontii* (Figure 1).

Our study likely only sampled millipedes from one of four known clades (LC3, Brewer et al. 2012), though we did not confirm this with sequencing. *Brachycybe lecontii* is a genetically diverse species with four major clades, which may represent cryptic species (Brewer et al. 2012). Future studies should sample from the remaining three clades to better understand if conserved OTUs simply reflect oversampling in a portion of *B. lecontii*’s range, or if these trends continue to point towards a possible mutualism.

It also remains to be seen if *B. lecontii* exhibit any choice or control over the fungi that live on their dead wood substrate. Two studies describing a novel animal-fungal symbiosis have explored similar questions, with red-cockaded woodpeckers in coastal North Carolina (Jusino et al. 2015). In the first study, researchers set out to determine if woodpeckers choose pines to excavate that already have specific fungi that make excavation easier (the tree selection hypothesis), or if they choose any pine, and subsequently alter the extant community by allowing ingress for their preferred fungi (the bird facilitation hypothesis). While these hypotheses could not be conclusively tested, the researchers found some evidence for both explanations. A follow-up study demonstrated that these birds actually facilitate fungal dispersal, providing conclusive evidence for the bird facilitation hypothesis (Jusino et al. 2016). It remains to be seen if *B. lecontii* have similar effects on their associated fungi and wood substrates.

## Conclusions

Studies on animal microbiomes have grown exponentially over the last decade, providing unique insights into the complex interactions that drive their lifestyles, behaviors, and biology. This study represents the first-ever mycobiome for the diverse and understudied Colobognatha, and one of the first using advanced methods with any millipede. *Brachycybe lecontii* associate with diverse fungal communities and are likely generalist fungal feeders, with different diets depending on location. We documented 2,863 fungal OTUs spanning 9 phyla, 100 orders, and 620 genera. More than half of the fungi in this study could not be assigned to an ecological guild, which may reflect novel ecological niches not captured in previous studies. Though this study found no evidence of a conserved fungal community, we did identify several OTUs that may be mutualists. Comparisons among other Colobognaths from the order Platydesmida, along with the three other fungus-feeding orders, are vital to identifying core fungal partners if they exist. Because our current knowledge of the Colobognatha and of fungal taxonomic and functional diversity is so fragmentary, so too is our understanding of how these organisms interact with each other and the surrounding environment.

## Acknowledgements

This work was supported by the following funding agencies. M.T.K. and A.M.M. were supported by WVU RSA Grant 1745 and National Geographic Grant NGS-74229R-20. A.M.M. received additional support from the West Virginia University Ruby Distinguished Doctoral Fellows Program. B.L. was supported by the USDA (USDA-ARS Project 8062-22410-007-000D). We thank Cameron Wilson for her assistance with geospatial analysis. All scientific collections were conducted with written permission by various public agencies. Sampling at Cove Lake State Park was conducted under permits held by Dr. Ernest Bernard & Gary Phillips with permission.

## Figure Captions

**Supplemental Figure 1:**
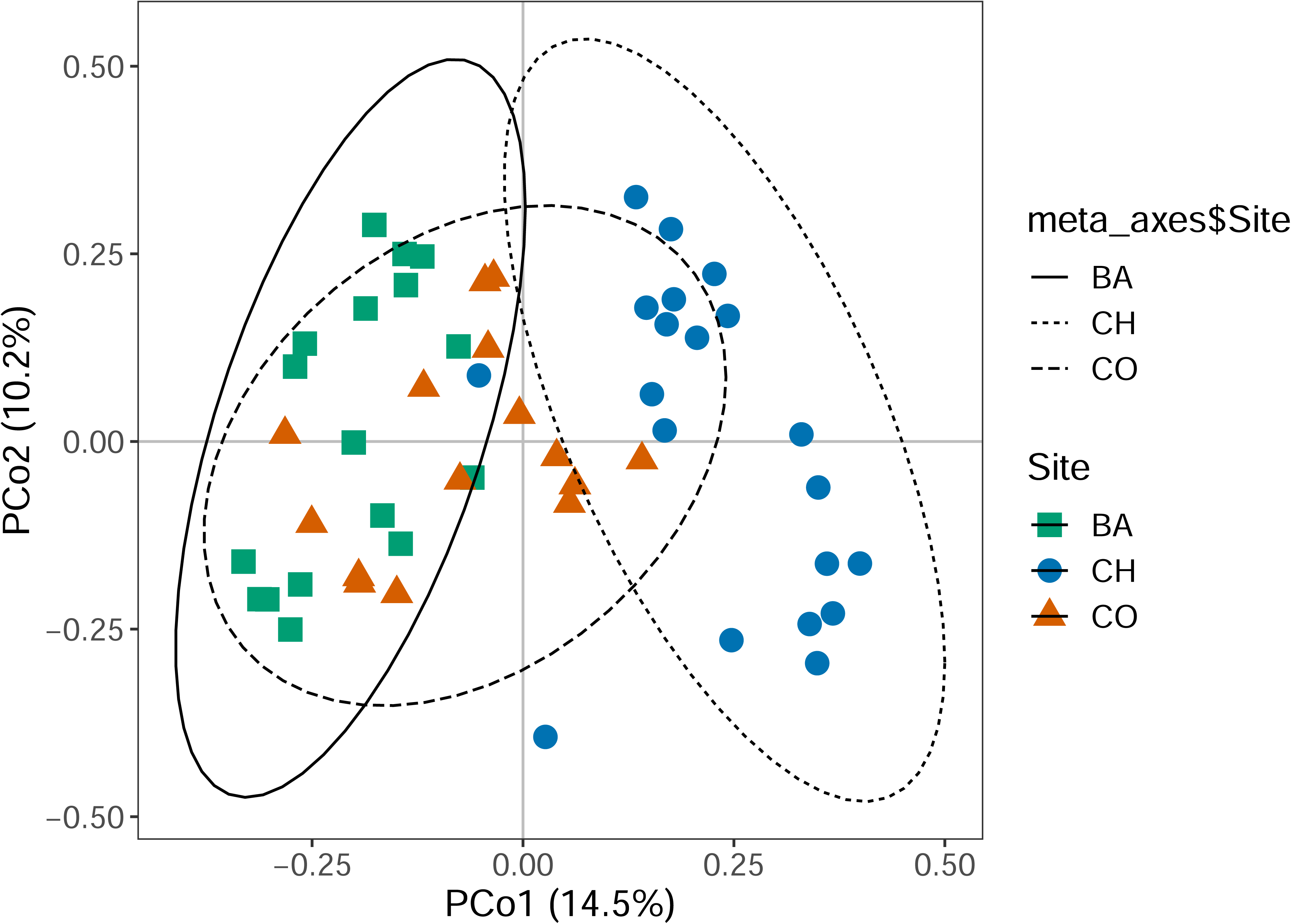
Principal coordinate analysis (PCoA) ordination plot showing Bray-Curtis based beta diversity for the culture-free dataset. Point shape & color indicates site: green squares = BA, blue circles = CH, and orange triangles = CO. Ellipses indicate 95% confidence intervals based on standard error of site centroids. Ellipse line type also indicates site: solid = BA, dotted = CH, dashed = CO.

**Supplemental Figure 2:**
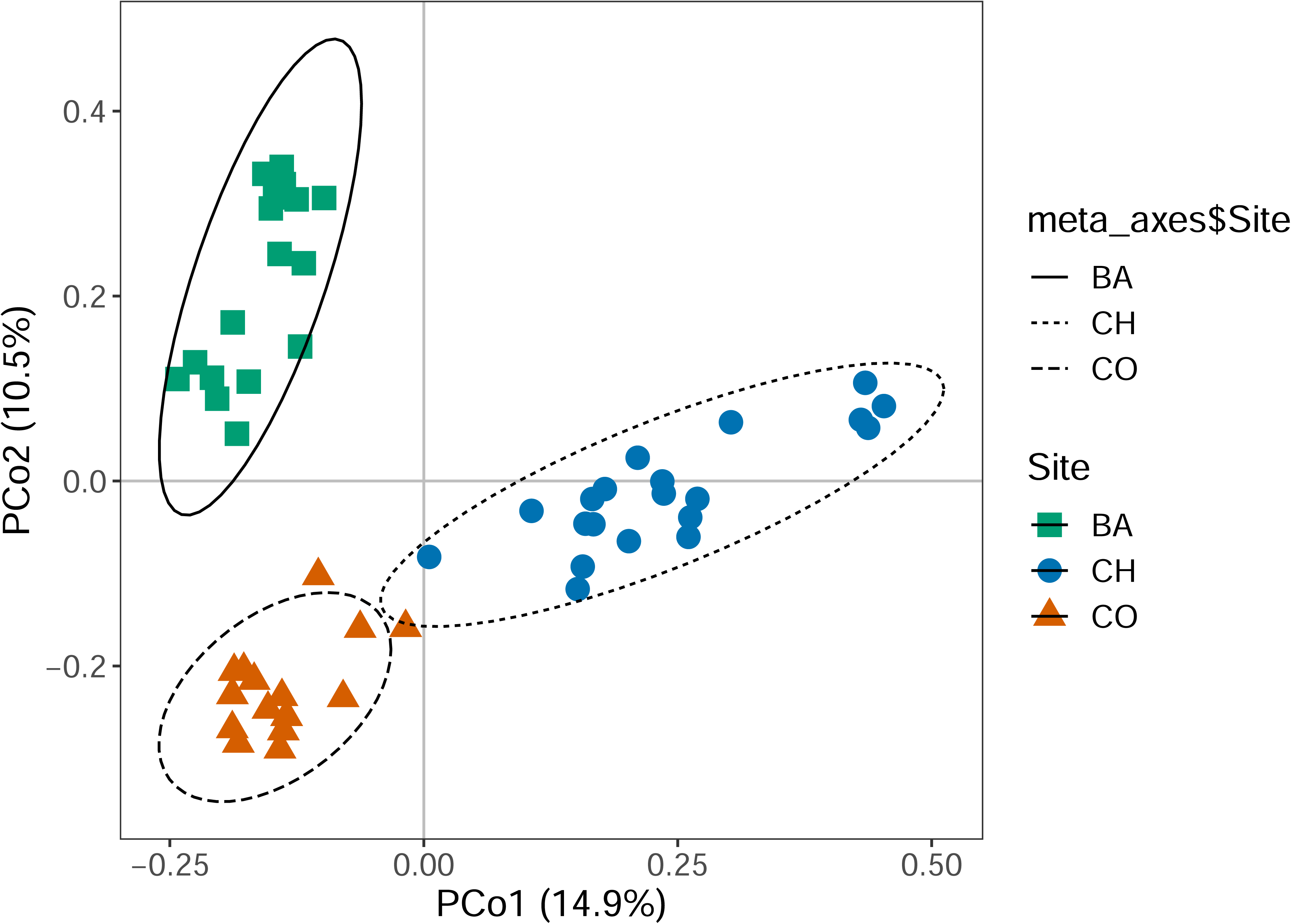
Principal coordinate analysis (PCoA) ordination plot showing βsim-based beta diversity for the culture-free dataset. Point shape & color indicates site: green squares = BA, blue circles = CH, and orange triangles = CO. Ellipses indicate 95% confidence intervals based on standard error of site centroids. Ellipse line type indicates site: solid = BA, dotted = CH, dashed = CO.

**Supplemental Figure 3:**
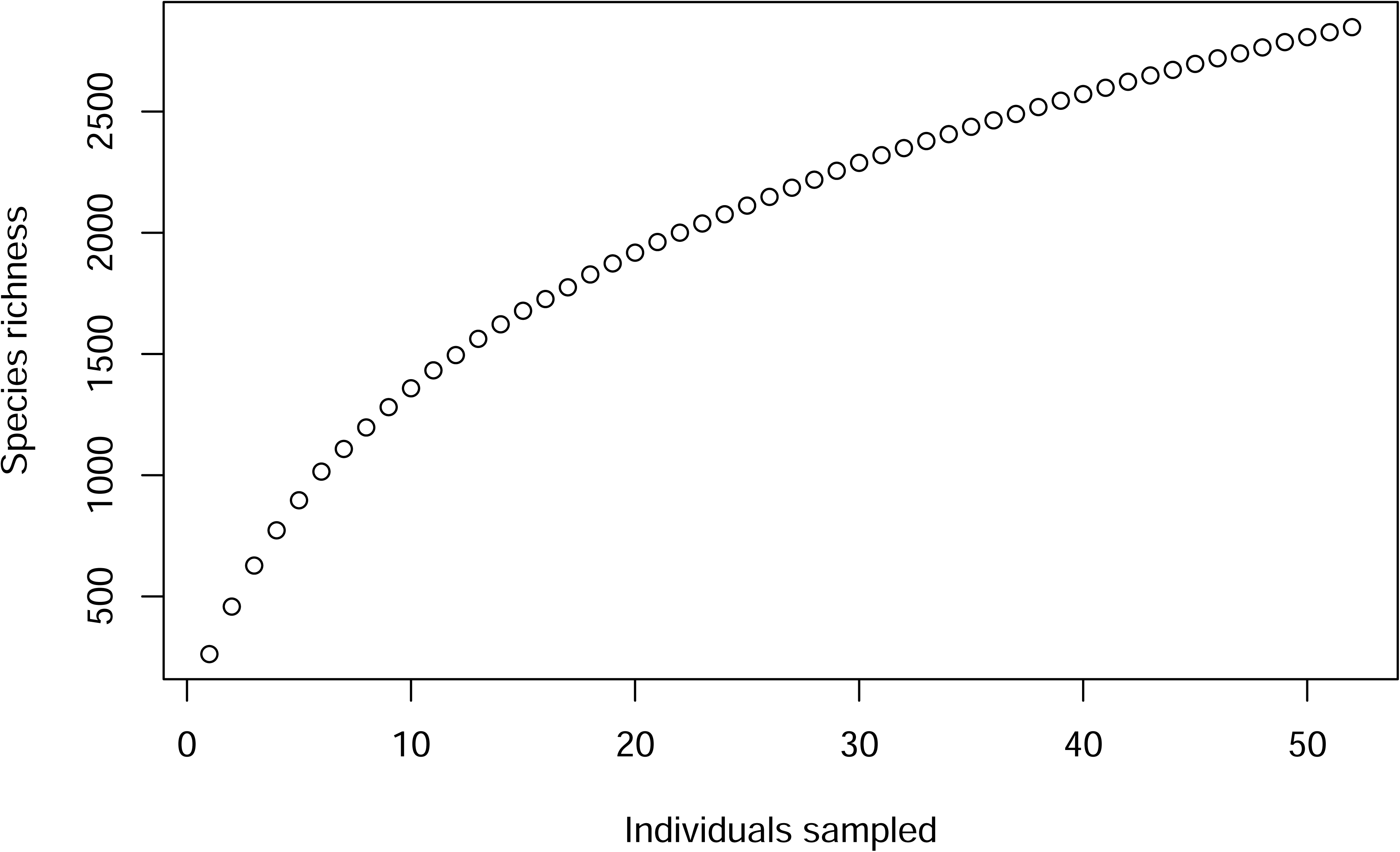
Taxon accumulation curve generated from random permutations of the culture-free fungal community data.

**Supplemental Figure 4:**
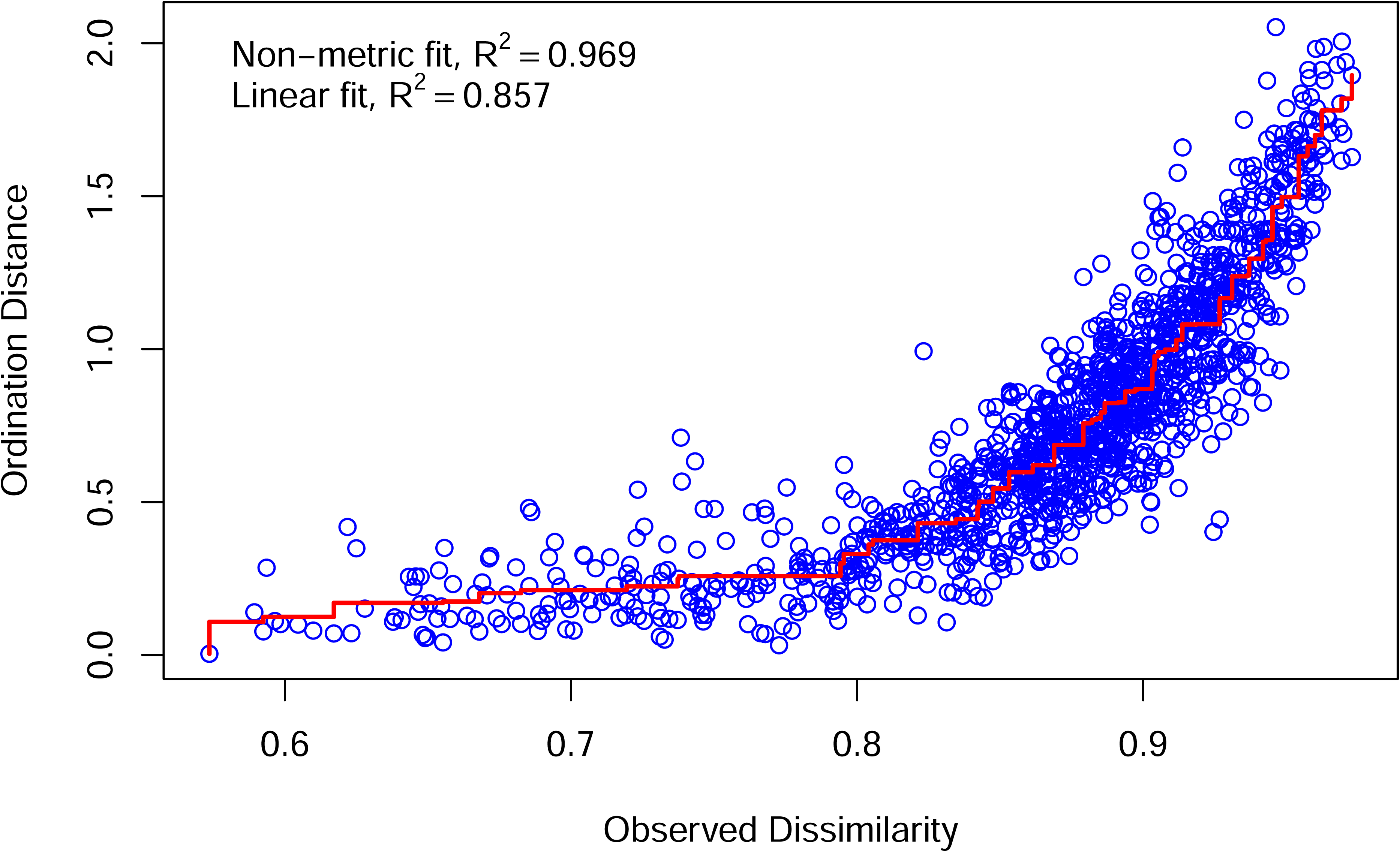
Shepard diagram showing ordination distances for the Bray-Curtis based NMDS shown in Figure 3.

**Supplemental Figure 5:**
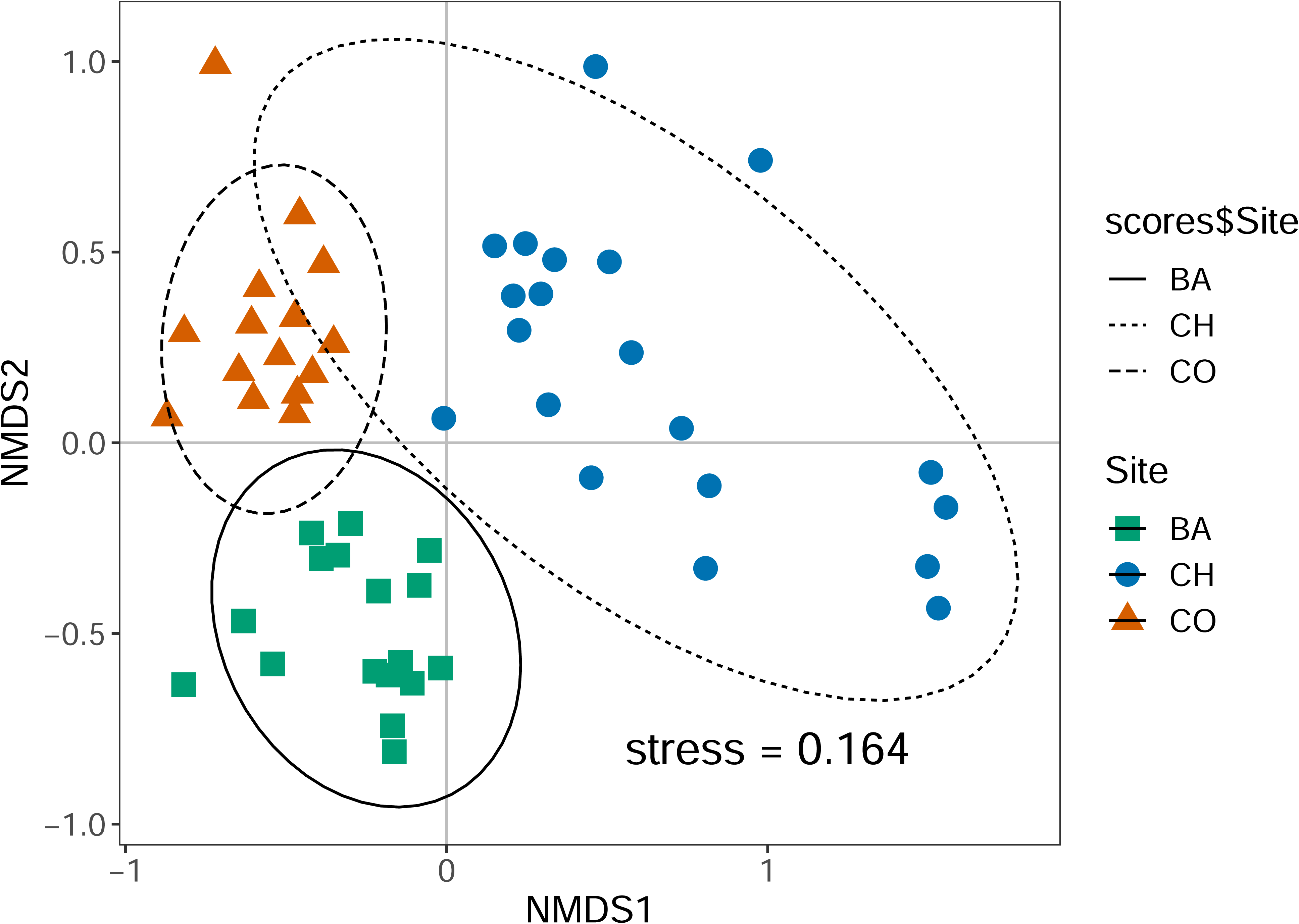
Nonmetric multidimensional scaling (NMDS) ordination plot showing βsim dissimilarity scores for all fungi recovered in the culture-free dataset. Point shape & color indicates site: green square = BA, blue circle = CH, orange triangle = CO. Ellipses indicate 95% confidence intervals based on standard error of site centroids. Ellipse line type also indicates site: solid = BA, dotted = CH, dashed = CO. Companion Shepard plot in Supp Fig. 6.

**Supplemental Figure 6:**
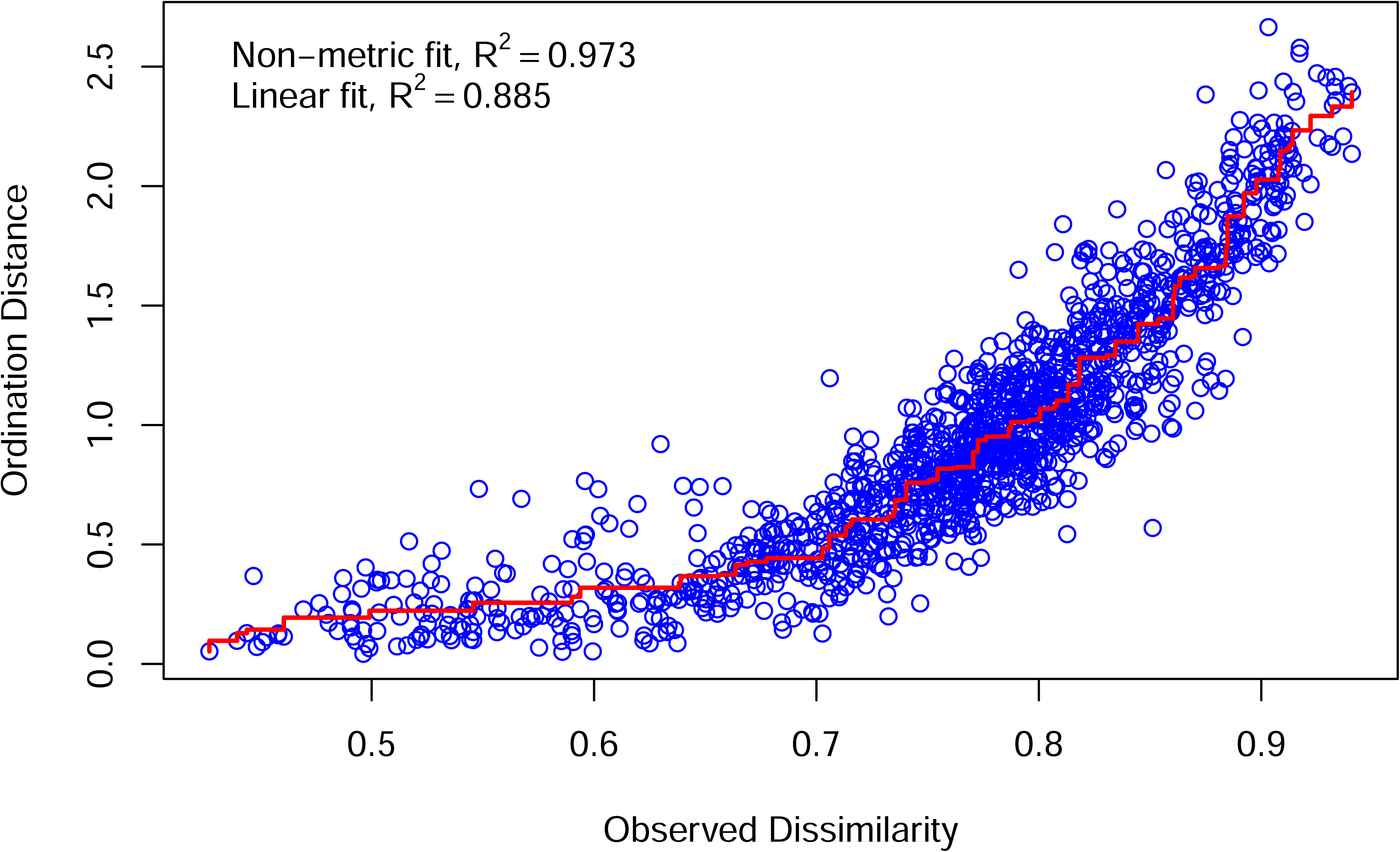
Shepard diagram showing ordination distances for the Bsim-based NMDS shown in Supplemental Figure 5.

**Supplemental Figure 7:**
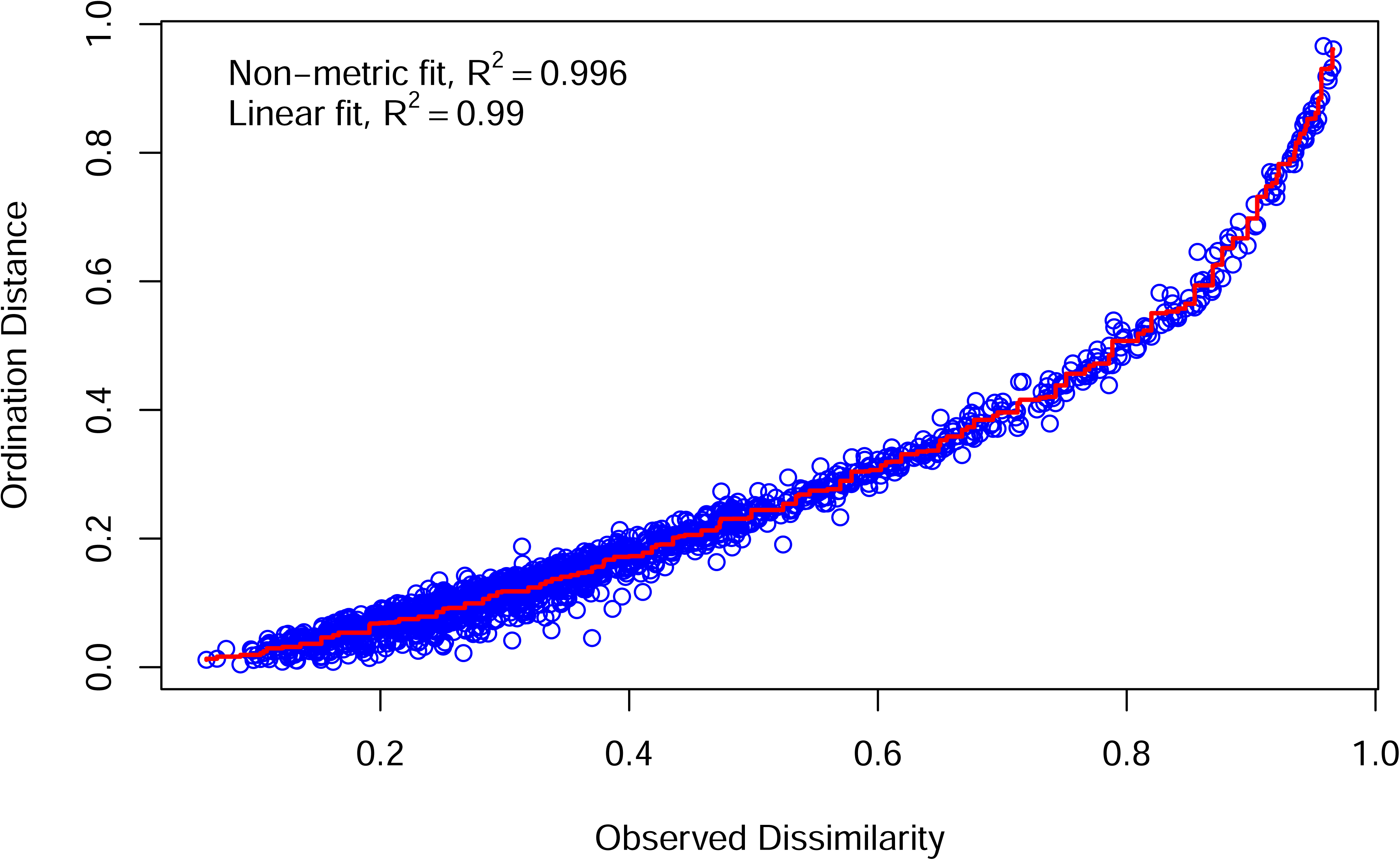
Shepard diagram showing ordination distances for the Bray-Curtis FunGuild NMDS shown in Figure 5.

## References

Aanen, D.K., de Fine Licht, H.H., Debets, A.J., Kerstes, N.A., Hoekstra, R.F. and Boomsma, J.J., 2009. High symbiont relatedness stabilizes mutualistic cooperation in fungus-growing termites. Science, 326(5956), pp.1103–1106.

Anderson, M.J., 2008. A new method for non-parametric multivariate analysis of variance. Austral Ecology, 26(1), pp.32–46.

Arbizu, P.M., 2017. *pairwiseAdonis*: Pairwise multilevel comparison using Adonis. R package version 0.4.

Banks, P., Funkhouser, E.M., Macias, A.M., Lovett, B., Meador, S., Hatch, A., Garraffo, H.M., Cartwright, K.C., Kasson, M.T., Marek, P.E. and Jones, T.H., 2024. The chemistry of the defensive secretions of three species of millipedes in the genus *Brachycybe*. Journal of Chemical Ecology, pp.1–11.

Bano, K., Bagyaraj, D.J. and Krishnamoorthy, R.V., 1976. Feeding activity of the millipede, Jonespeltis splendidus Verhoeff and soil humification. Proceedings of the Indian Academy of Sciences-Section B, 83, pp.1–11.

Bartlett, M.S., 1937. Properties of sufficiency and statistical tests. Proceedings of the Royal Society of London. Series A-Mathematical and Physical Sciences, 160(901), pp.268–282.

Beaver, R.A, 1989. Insect-fungus relationships in the bark and ambrosia beetles. In: Wilding, N., et al. (eds.). “Insect-Fungus Interactions.” Academic Press, Ltd. pp.121–144.

von Beeren, C., Mair, M. M., and Witte, V. 2014. Discovery of a second mushroom-harvesting ant (Hymenoptera: Formicidae) in Malayan tropical rainforests. Myrmecological News, 20, pp.37–42.

Ben-Shachar, M.S., Lüdecke, D., and Makowski, D., 2020. effectsize: Estimation of effect size indices and standardized parameters. Journal of Open Source Software, 5(56), pp.2815.

Bidartondo, M.I., and Gardes, M, 2005. Fungal diversity in molecular terms: profiling, identification, and quantification in the environment. In: Dighton, J., et al. (eds.). “The Fungal Community: Its Organization and Role in the Ecosystem.” CRC Press, Taylor & Francis Group. pp.215–239.

Biedermann, P.H., and Vega, F.E., 2020. Ecology and evolution of insect–fungus mutualisms. Annual Eeview of Entomology, 65(1), pp.431–455.

Birkemoe, T., Jacobsen, R.M., Sverdrup-Thygeson, A., Biedermann, P.H.W., 2018. Insect-Fungus Interactions in Dead Wood Systems. In: Ulyshen, M. (ed.). “Saproxylic Insects.” Zoological Monographs, vol 1. Springer. pp.377–427.

Brewer, M.S., Spruill, C.L., Rao, N.S. and Bond, J.E., 2012. Phylogenetics of the millipede genus *Brachycybe* Wood, 1864 (Diplopoda: Platydesmida: Andrognathidae): Patterns of deep evolutionary history and recent speciation. Molecular Phylogenetics and Evolution, 64(1), pp.232–242.

BugGuide. Available from https://bugguide.net/.

Byzov, B., 2006. Intestinal microbiota of millipedes. In: König, H., and Varma, A. (eds.). “Intestinal Microorganisms of Termites and Other Invertebrates.” Springer Berlin, Heidelberg. pp.89–114.

Byzov, B.A., Thanh, V.N. and Babjeva, I.P., 1993a. Interrelationships between yeasts and soil diplopods. Soil Biology and Biochemistry, 25(8), pp.1119–1126.

Byzov, B.A., Thanh, V.N. and Babjeva, I.P., 1993b. Yeasts associated with soil invertebrates. Biology and Fertility of Soils, 16, pp.183–187.

Callahan, B.J., McMurdie, P.J., Rosen, M.J., Han, A.W., Johnson, A.J.A. and Holmes, S.P., 2016. DADA2: High-resolution sample inference from Illumina amplicon data. Nature Methods, 13(7), pp.581–583.

Camacho, C., Coulouris, G., Avagyan, V., Ma, N., Papadopoulos, J., Bealer, K., & Madden, T.L., 2009. BLAST+: architecture and applications. BMC Bioinformatics, 10, p.421.

Carreiro, S.C., Pagnocca, F.C., Bacci Jr., M., Bueno, O.C., Hebling, M.J.A., and Middelhoven, W.J., 2002. Occurrence of killer yeasts in leaf-cutting ant nests. Folia Microbiologica, 47(3) pp.259–262.

Claridge, A.W. and Trappe, J.M., 2005. Sporocarp mycophagy: nutritional, behavioral, evolutionary, and physiological aspects. In: Dighton, J. et al. (eds). “The Fungal Community: Its Organization and Role in the Ecosytem.” 3rd edn. Taylor & Francis Group. pp.599–611.

Coutts, M.P., and Dolezal, J.E., 1969. Emplacement of fungal spores by the woodwasp, *Sirex noctilio*, during oviposition. Forest Science, 15(4), pp.412–416.

Davis, T.S., Hofstetter, R.W., Foster, J.T., Foote, N.E. and Keim, P., 2011. Interactions between the yeast *Ogataea pini* and filamentous fungi associated with the western pine beetle. Microbial Ecology, 61, pp.626–634.

Dexter, E., Rollwagen-Bollens, G. and Bollens, S.M., 2018. The trouble with stress: A flexible method for the evaluation of nonmetric multidimensional scaling. Limnology and Oceanography: Methods, 16(7), pp.434–443.

Dhivya, A. and Alagesan, P., 2018. Isolation and identification of microbial load in the gut and faeces of millipede *Spinotarsus colosseus*. World Journal of Zoology, 13(2), pp.4–9.

Douglas, A.E., 2022. “Insects and Their Beneficial Microbes.” Princeton University Press.

Elliott, T.F., Bower, D.S., and Vernes, K., 2019a. Reptilian mycophagy: A global review of mutually beneficial associations between reptiles and macrofungi. Mycosphere, 10(1), pp.776–797.

Elliott, T.F., Jusino, M.A., Trappe, J.M., Lepp, H., Ballard, G.A., Bruhl, J.J. and Vernes, K., 2019b. A global review of the ecological significance of symbiotic associations between birds and fungi. Fungal Diversity, 98(1), pp.161–194.

Elliott, T.F., Truong, C., Jackson, S.M., Zúñiga, C.L., Trappe, J.M. and Vernes, K., 2022. Mammalian mycophagy: a global review of ecosystem interactions between mammals and fungi. Fungal Systematics and Evolution, 9(1), pp.99–159.

Fisher, R.A., Corbet, A.S. and Williams, C.B., 1943. The relation between the number of species and the number of individuals in a random sample of an animal population. The Journal of Animal Ecology, 12(1), pp.42–58.

Flórez, L.V., Biedermann, P.H., Engl, T., and Kaltenpoth, M., 2015. Defensive symbioses of animals with prokaryotic and eukaryotic microorganisms. Natural Product Reports, 32(7), pp.904–936.

Fontanetti, C.S., Moreira-de-Sousa, C., Pinheiro, T.G., de Souza, R.B., and Francisco, A, 2015. Diplopoda—digestive system. In: Minelli, A. (ed.). “The Myriapoda”, vol. 2, in series “Treatise of Zoology - Anatomy, Taxonomy, and Biology” Koninklijke Brill NV. pp. 109–128.

Frøslev, T.G., Kjøller, R., Bruun, H.H., Ejrnæs, R., Brunbjerg, A.K., Pietroni, C. and Hansen, A.J., 2017. Algorithm for post-clustering curation of DNA amplicon data yields reliable biodiversity estimates. Nature Communications, 8(1), p.1188.

Games, P.A. and Howell, J.F., 1976. Pairwise multiple comparison procedures with unequal n’s and/or variances: a Monte Carlo study. Journal of Educational Statistics, 1(2), pp.113–125.

Geli-Cruz, O.J., Santos-Flores, C.J., Cafaro, M.J., Ropelewski, A. and Van Dam, A.R., 2023. Benchmarking assembly free nanopore read mappers to classify complex millipede gut microbiota via Oxford Nanopore Sequencing Technology. Journal of Biological Methods, 10.

Gilliam, M., Taber III, S., Lorenz, B.J,. and Prest, D.B., 1988. Factors affecting development of chalkbrood disease in colonies of honey bees, *Apis mellifera*, fed pollen contaminated with *Ascosphaera apis*. Journal of Invertebrate Pathology, 52(2), pp.314–325.

Gomi, S., Imamura, K.I., Yaguchi, T., Kodama, Y., Minowa, N. and Koyama, M., 1994. PF1018, a novel insecticidal compound produced by *Humicola* sp. The Journal of Antibiotics, 47(5), pp.571–580.

Gower, J.C., 1966. Some distance properties of latent root and vector methods used in multivariate analysis. Biometrika, 53(3-4), pp.325–338.

Green, K., Tory, M.K., Mitchell, A.T., Tennant, P. and May, T.W., 1999. The diet of the long-footed potoroo (*Potorous longipes*). Australian Journal of Ecology, 24(2), pp.151–156.

Guo, W., Wang, W., Tang, J., Li, T. and Li, X., 2023. Genome analysis and genomic comparison of a fungal cultivar of the nonsocial weevil *Euops chinensis* reveals its plant decomposition and protective roles in fungus-farming mutualism. Frontiers in Microbiology, 14, pp.1048910.

Hajek, A.E., Morris, E.E., and Hendry, T.A., 2019. Context-dependent interactions of insects and defensive symbionts: insights from a novel system in siricid woodwasps. Current Opinion in Insect Science, 33, pp.77–83.

Hammond, P.M, and Lawrence, J.F., 1989. Mycophagy in insects: a summary. In: Wilding, N., et al. (eds.). “Insect-Fungus Interactions.” Academic Press, Ltd. pp.275–323.

Hanski, I., 1989. Fungivory: fungi, insects and ecology. In: Wilding, N., et al. (eds.). “Insect-Fungus Interactions.” Academic Press, Ltd. pp.25–68.

Heděnec, P., Zheng, H., Siqueira, D.P., Peng, Y., Schmidt, I.K., Frøslev, T.G., Kjøller, R., Li, H., Frouz, J. and Vesterdal, L., 2023. Litter chemistry of common European tree species drives the feeding preference and consumption rate of soil invertebrates, and shapes the diversity and structure of gut and faecal microbiomes. Soil Biology and Biochemistry, 177, p.108918.

Hervey, A. and Nair, M.S.R., 1979. Antibiotic metabolite of a fungus cultivated by gardening ants. Mycologia, 71(5), pp.1064–1066.

Hopkin, S.P., and Read, H.J. 1992. “The Biology of Millipedes.” Oxford University Press. pp.43–60.

Hopple Jr, J.S. and Vilgalys, R., 1999. Phylogenetic relationships in the mushroom genus *Coprinus* and dark-spored allies based on sequence data from the nuclear gene coding for the large ribosomal subunit RNA: divergent domains, outgroups, and monophyly. Molecular Phylogenetics and Evolution, 13(1), pp.1–19.

Hulcr, J., and Stelinski, L.L., 2017. The ambrosia symbiosis: from evolutionary ecology to practical management. Annual Review of Entomology, 62, pp.285–303.

iNaturalist. Available from https://www.inaturalist.org.

IUCN. 2024. The IUCN Red List of Threatened Species. Version 2024-1. https://www.iucnredlist.org. Accessed on 10 July 2024.

Jarosz, J. and Kania, G., 2000. The question of whether gut microflora of the millipede *Ommatoiulus sabulosus* could function as a threshold to food infections. Pedobiologia, 44(6), pp.705–708.

Jones, T.H., Harrison, D.P., Menegatti, C., Mevers, E., Knott, K., Marek, P., Hennen, D.A., Kasson, M.T., Macias, A.M., Lovett, B. and Saporito, R.A., 2022. Deoxybuzonamine isomers from the millipede *Brachycybe lecontii* (Platydesmida: Andrognathidae). Journal of Natural Products, 85(4), pp.1134–1140.

Jusino, M.A., Lindner, D.L., Banik, M.T., Rose, K.R. and Walters, J.R., 2016. Experimental evidence of a symbiosis between red-cockaded woodpeckers and fungi. Proceedings of the Royal Society B: Biological Sciences, 283(1827), pp.20160106.

Jusino, M.A., Lindner, D.L., Banik, M.T. and Walters, J.R., 2015. Heart rot hotel: fungal communities in red-cockaded woodpecker excavations. Fungal Ecology, 14, pp.33–43.

Kahlke, T. and Ralph, P.J., 2019. BASTA–Taxonomic classification of sequences and sequence bins using last common ancestor estimations. Methods in Ecology and Evolution, 10(1), pp.100–103.

Kassambara, A., 2023. *rstatix*: Pipe-friendly framework for basic statistical tests. R package version 0.7.2.

Knapp, B.A., Seeber, J., Podmirseg, S.M., Rief, A., Meyer, E. and Insam, H., 2009. Molecular fingerprinting analysis of the gut microbiota of *Cylindroiulus fulviceps* (Diplopoda). Pedobiologia, 52(5), pp.325–336.

Koubová, A., Lorenc, F., Horváthová, T., Chroňáková, A. and Šustr, V., 2023. Millipede gut-derived microbes as a potential source of cellulolytic enzymes. World Journal of Microbiology and Biotechnology, 39(7), pp.169.

Kudo, S., Akagi, Y., Hiraoka, S., Tanabe, T., and Morimoto, G., 2010. Exclusive male egg care and determinants of brooding success in a millipede. Ethology, 117(1), pp.19–27.

Lehmkuhl, J.F., Gould, L.E., Cázares, E. and Hosford, D.R., 2004. Truffle abundance and mycophagy by northern flying squirrels in eastern Washington forests. Forest Ecology and Management, 200(1-3), pp.49–65.

Levene, H., 1960. Robust tests for equality of variances. In: Olkin, I., et al. (eds.). “Contributions to Probability and Statistics: Essays in Honor of Harold Hotelling.*”* Stanford University Press. pp.278–292.

Lewis, J.G.E., 1971. The life history and ecology of three paradoxosomatid millipedes (Diplopoda: Polydesmida) in northern Nigeria. Journal of Zoology, 165(4), pp.431–452.

Lücking, R., Aime, M.C., Robbertse, B., Miller, A.N., Ariyawansa, H.A., Aoki, T., Cardinali, G., Crous, P.W., Druzhinina, I.S., Geiser, D.M. and Hawksworth, D.L., 2020. Unambiguous identification of fungi: where do we stand and how accurate and precise is fungal DNA barcoding?. IMA Fungus, 11(1), pp.14.

Macias, A.M., Marek, P.E., Morrissey, E.M., Brewer, M.S., Short, D.P., Stauder, C.M., Wickert, K.L., Berger, M.C., Metheny, A.M., Stajich, J.E. and Boyce, G., 2019. Diversity and function of fungi associated with the fungivorous millipede, *Brachycybe lecontii*. Fungal Ecology, 41, pp.187–197.

Maser, C. and Maser, Z., 1988. Mycophagy of red-backed voles, Clethrionomys californicus and C. gapperi. The Great Basin Naturalist, 48(2), pp.269–273.

Martin, M.M., 1979. Biochemical implications of insect mycophagy. Biological Reviews, 54, pp.1–21.

Menezes, C., Vollett-Neto, A., Marsaioli, A.J., Zampirei, D., Fontoura, I.C., Luchessi, A.D., and Imperatriz-Fonseca, V.L., 2015. A Brazilian social bee must cultivate fungus to survive. Current Biology, 25(21), pp.2851–2855.

MilliBase. Available from https://www.millibase.org.

Moritz, L., Blanke, A., Hammel, J.U., and Wesener, T., 2021. First steps toward suctorial feeding in millipedes: comparative morphology of the head of the Platydesmida (Diplopoda: Colobognatha). Invertebrate Biology, 140(2), e12312.

Moritz, L., Borisova, E., Hammel, J.U., Blanke, A., and Wesener, T., 2022. A previously unknown feeding mode in millipedes and the convergence of fluid feeding across arthropods. Science Advances, 8(7), eabm057.

Mueller, U.G., Gerardo, N.M., Aanen, D.K., Six, D.L., and Schultz, T.R., 2005. The evolution of agriculture in insects. Annual Review of Ecology, Evolution, and Systematics, 36, pp.563–595.

Murakami, Y., 1962. Postembryonic development of the common Myriapoda of Japan. XII. Life history of *Bazillozonium nodulosum* (Colobognatha, Platydesmidae). Zoological Magazine, 72, pp.40–47.

Mwabvu, T., Mswaka, A. and Mlambo, G., 2003. A preliminary survey of micro-organisms in the gut and pellets of a tropical millipede *Doratogonus uncinatus* Attems (Diplopoda, Spirostreptida, Spirostreptidae). Global Journal of Pure and Applied Sciences, 9(2), pp.171–176.

Nakashima, T., Izawa, T., Ogura, K., Maeda, M. and Tanaka, T., 1982. Isolation of some microorganisms associated with five species of ambrosia beetles and two kinds of antibiotics produced by Xv-3 strain in these isolates. *Journal of the Faculty of Agriculture*, Hokkaido University, 61(1), pp.60–72.

Neal, B.P., Honisch, B., Warrender, T., Williams, G.J., Work, T.M. and Price, N.N., 2020. Possible control of acute outbreaks of a marine fungal pathogen by nominally herbivorous tropical reef fish. Oecologia, 193(3), pp.603–617.

Neu, A.T., Allen, E.E. and Roy, K., 2021. Defining and quantifying the core microbiome: challenges and prospects. Proceedings of the National Academy of Sciences, 118(51), p.e2104429118.

Nguyen, N.H., Song, Z., Bates, S.T., Branco, S., Tedersoo, L., Menke, J., Schilling, J.S. and Kennedy, P.G., 2016. FUNGuild: an open annotation tool for parsing fungal community datasets by ecological guild. Fungal Ecology, 20, pp.241–248.

Nishino, T., Mukai, H., Moriyama, M., Hosokawa, T., Tanahashi, M., Tachikawa, S., Nikoh, N., Koga, R., and Fukatsu, T., 2024. Defensive fungal symbiosis on insect hindlegs. BioRxiv. https://www.biorxiv.org/content/10.1101/2024.03.25.586038v1

Nweze, J.E., Šustr, V., Brune, A. and Angel, R., 2024. Functional similarity, despite taxonomical divergence in the millipede gut microbiota, points to a common trophic strategy. Microbiome, 12(1), pp.16.

Oksanen, J., Blanchet, F.G., Friendly, M., Kindt, R., Legendre, P., McGlinn, D., Minchin, P.R., O’Hara, R.B., Simpson, G.L., Solymos, P., Stevens, M.H., Szoecs, E., and Wagner, H., 2020. *vegan*: Community ecology package. R package version 2.5–7.

Ori, F., Menotta, M., Leonardi, M., Amicucci, A., Zambonelli, A., Covès, H., Selosse, M.A., Schneider-Maunoury, L., Pacioni, G. and Iotti, M., 2021. Effect of slug mycophagy on *Tuber aestivum* spores. Fungal Biology, 125(10), pp.796–805.

Palmer, J.M., Jusino, M.A., Banik, M.T. and Lindner, D.L., 2018. Non-biological synthetic spike-in controls and the AMPtk software pipeline improve mycobiome data. PeerJ, 6, p.e4925.

Pan, J., Bhardwaj, M., Faulkner, J.R., Nagabhyru, P., Charlton, N.D., Higashi, R.M., Miller, A.F., Young, C.A., Grossman, R.B. and Schardl, C.L., 2014. Ether bridge formation in loline alkaloid biosynthesis. Phytochemistry, 98, pp.60–68.

Pearson, K., 1895. Notes on regression and inheritance in the case of two parents. Proceedings of the Royal Society of London, 58, pp.240–242.

Pölme, S., Abarenkov, K., Henrik Nilsson, R., Lindahl, B.D., Clemmensen, K.E., Kauserud, H., Nguyen, N., Kjøller, R., Bates, S.T., Baldrian, P. and Frøslev, T.G., 2020. FungalTraits: a user-friendly traits database of fungi and fungus-like stramenopiles. Fungal Diversity, 105, pp.1–16.

R Core Team, 2020. R: A language and environment for statistical computing. R Foundation for Statistical Computing, Vienna, Austria. Available from https://www.R-project.org/.

Read, H.J., and Enghoff, H., 2009. The order Siphonophorida – A taxonomist’s nightmare? Lessons from a Brazilian collection. Soil Organisms, 81(3), pp.543–556.

Reynolds, N.K., Benny, G.L., Ho, H.M., Hou, Y.H., Crous, P.W. and Smith, M.E., 2019. Phylogenetic and morphological analyses of the mycoparasitic genus *Piptocephalis*. Mycologia, 111(1), pp.54–68.

Reynolds, N.K., Jusino, M.A., Stajich, J.E. and Smith, M.E., 2022. Understudied, underrepresented, and unknown: methodological biases that limit detection of early diverging fungi from environmental samples. Molecular Ecology Resources, 22(3), pp.1065–1085.

Rodrigues, A., Cable, R.N., Mueller, U.G., Bacci, M. and Pagnocca, F.C., 2009. Antagonistic interactions between garden yeasts and microfungal garden pathogens of leaf-cutting ants. Antonie van Leeuwenhoek, 96, pp.331–342.

Rohlfs, M., and Kürschner, L., 2010. Saprophagous insect larvae, *Drosophila melanogaster*, profit from increased species richness in beneficial microbes. Journal of Applied Entomology, 134(8), pp.667–671.

Rossa-Feres, D.D.C., Jim, J. and Fonseca, M.G., 2004. Diets of tadpoles from a temporary pond in southeastern Brazil (Amphibia, Anura). Revista Brasileira de Zoologia, 21, pp.745–754.

Ruess, L., and Lussenhop, M., 2005.Trophic interactions of fungi and animals. In: Dighton, J., et al. (eds.). “The Fungal Community: Its Organization and Role in the Ecosystem.” CRC Press, Taylor & Francis Group. pp.581–599.

Santamaria, B., Verbeken, A. and Haelewaters, D., 2023. Mycophagy: A global review of interactions between invertebrates and fungi. Journal of Fungi, 9(2).

Sardar, P., Šustr, V., Chroňáková, A., Lorenc, F. and Faktorová, L., 2022. De novo metatranscriptomic exploration of gene function in the millipede holobiont. Scientific Reports, 12(1), pp.16173.

Schoch, C.L., Ciufo, S., Domrachev, M., Hotton, C.L., Kannan, S., Khovanskaya, R., Leipe, D., Mcveigh, R., O’Neill, K., Robbertse, B. Sharma, S., Soussov, V., Sullivan, J.P., Sun, L., Turner, S., and Karsch-Mizrachi, I., 2020. NCBI Taxonomy: a comprehensive update on curation, resources and tools. Database, 2020, pp.1–21.

Schoch, C.L., Seifert, K.A., Huhndorf, S., Robert, V., Spouge, J.L., Levesque, C.A., Chen, W., Fungal Barcoding Consortium, Fungal Barcoding Consortium Author List, Bolchacova, E. and Voigt, K., 2012. Nuclear ribosomal internal transcribed spacer (ITS) region as a universal DNA barcode marker for Fungi. Proceedings of the National Academy of Sciences, 109(16), pp.6241–6246.

Shannon, C.E., 1948. A mathematical theory of communication. The Bell System Technical Journal, 27, pp.379–656.

Shapiro, S.S. and Wilk, M.B., 1965. An analysis of variance test for normality (complete samples). Biometrika, 52(3-4), pp.591–611.

Shear, W.A., 2015. The chemical defenses of millipedes (Diplopoda): biochemistry, physiology and ecology. Biochemical Systematics and Ecology 61, pp.78–117.

Shepard, R.N., 1962a. The analysis of proximities: Multidimensional scaling with an unknown distance function. I. Psychometrika, 27, pp.125–140.

Shepard, R.N. ,1962b. The analysis of proximities: Multidimensional scaling with an unknown distance function. II. Psychometrika, 27, pp.219–246.

Shorter, P.L., Hennen, D.A., and Marek, P.E., 2018. Cryptic diversity in *Andrognathus corticarius* Cope, 1869 and description of a new *Andrognathus* species from New Mexico (Diplopoda, Platydesmida, Andrognathidae). ZooKeys, 786, pp.19–41.

Shukla, S.P., Plata, C., Reichelt, M., Steiger, S., Heckel, D.G., Kaltenpoth, M., Vilcinskas, A., and Vogel, H., 2018. Microbiome-assisted carrion preservation aids larval development in a burying beetle. Proceedings of the National Academy of Sciences, 115(44), pp.11274–11279.

Silliman, B.R., and Newell, S.Y., 2003. Fungal farming in a snail. Proceedings of the National Academy of Sciences, 100(26), pp.15643–15648.

Sinclair, E.A., Danks, A., and Wayne, A.F., 1996. Rediscovery of Gilbert’s potoroo, *Potorous tridactylus*. Australian Mammalogy Notes, 19(1), pp.69–72.

Taylor, E.C., 1982. Fungal preference by a desert millipede, *Orthoporus ornatus* (Spirostreptidae). Pedobiologia, 23(5), pp.331–336.

Thanh, V.N., Byzov, B.A. and Babeva, I.P., 1994. Sensitivity of yeasts to digestive fluid in the gut of soil diplopods *Pachyiulus flavipes* CL Koch. Microbiology, 63, pp.406–409.

Thomas, P.W. and Thomas, H.W., 2022. Mycorrhizal fungi and invertebrates: impacts on *Tuber melanosporum* ascospore dispersal and lifecycle by isopod mycophagy. Food Webs, 33, e00260.

Toki, W., Tanahashi, M., Togashi, K. and Fukatsu, T., 2012. Fungal farming in a non-social beetle. PLoS ONE, 7(7), e41893.

USFWS ECOS (United States Fish and Wildlife Service: Environmental Conservation Online Service). Accessed August 2024. *Available from:* https://ecos.fws.gov/ecp/

Van Bael, S.A., Fernández-Marín, H., Valencia, M.C., Rojas, E.I., Wcislo, W.T. and Herre, E.A., 2009. Two fungal symbioses collide: endophytic fungi are not welcome in leaf-cutting ant gardens. Proceedings of the Royal Society B: Biological Sciences, 276(1666), pp.2419–2426.

Wang, Y., Mueller, U.G. and Clardy, J.O.N., 1999. Antifungal diketopiperazines from symbiotic fungus of fungus-growing ant *Cyphomyrmex minutus*. Journal of Chemical Ecology, 25, pp.935–941.

Welch, B.L., 1951. On the comparison of several mean values: an alternative approach. Biometrika, 38(3-4), pp.330–336.

Wesener, T., and Moritz, L, 2018. Checklist of the Myriapoda in Cretaceous Burmese amber and a correction of the Myriapoda identified by Zhang (2017). Check List 14, pp.1131–1140.

White, T.J., 1990. Amplification and direct sequencing of fungal ribosomal RNA genes for phylogenetics. PCR Protocols: A guide to methods and applications/Academic Press, Inc.

Wickham, H., 2016. *ggplot2*: Elegant graphics for data analysis. Springer-Verlag New York. R package version 3.4.4.

Wilcoxon, F., 1992. Individual comparisons by ranking methods. In: “Breakthroughs in statistics: Methodology and distribution.” Springer New York. pp.196–202.

Wong, V.L., Hennen, D.A., Macias, A.M., Brewer, M.S., Kasson, M.T. and Marek, P., 2020. Natural history of the social millipede *Brachycybe lecontii* Wood, 1864. Biodiversity Data Journal, 8, p.e50770.

Wood, W., Hanke, F., Kubo, I., Carrol,l J., and Crews, P., 2000. Buzonamine, a new alkaloid from the defensive secretion of the millipede, *Buzonium crassipes*. Biochemical Systematics and Ecology, 28(4), pp.305–312.

Xu, K., Yuan, X.L., Li, C. and Li, X.D., 2020. Recent discovery of heterocyclic alkaloids from marine-derived *Aspergillus* species. Marine Drugs, 18(1), pp.54.

Zhang, Y., Li, S., Li, H., Wang, R., Zhang, K.Q. and Xu, J., 2020. Fungi-nematode interactions: diversity, ecology, and biocontrol prospects in agriculture. Journal of Fungi, 6(4).

